# Elf1 promotes Rad26’s interaction with lesion-arrested Pol II for transcription-coupled repair

**DOI:** 10.1101/2023.09.08.556893

**Authors:** Reta D Sarsam, Jun Xu, Indrajit Lahiri, Wenzhi Gong, Juntaek Oh, Zhen Zhou, Jenny Chong, Nan Hao, Shisheng Li, Dong Wang, Andres E. Leschziner

**Affiliations:** Department of Cellular & Molecular Medicine, University of California San Diego, La Jolla, CA 92093; Division of Pharmaceutical Sciences, Skaggs School of Pharmacy & Pharmaceutical Sciences, University of California San Diego, La Jolla, CA 92093; Department of Molecular Biology, School of Biological Sciences, University of California San Diego, La Jolla, CA 92093; Department of Chemistry and Biochemistry, University of California San Diego, La Jolla, CA 92093; Department of Comparative Biomedical Sciences, School of Veterinary Medicine, Louisiana State University, Baton Rouge, LA 70803; Charles River Laboratories, Durham, NC 27703; Genetics and Metabolism Department, The Children’s Hospital, School of Medicine, Zhejiang University, National Clinical Research Center for Child Health, Hangzhou 310052, China; School of Biosciences, University of Sheffield, UK; Department of Pharmacy, College of Pharmacy, Kyung Hee University; Seoul 02447, Republic of Korea

## Abstract

Transcription-coupled nucleotide excision repair (TC-NER) is a highly conserved DNA repair pathway that removes bulky lesions in the transcribed genome. Cockayne syndrome B protein (CSB), or its yeast ortholog Rad26, has been known for decades to play important roles in the lesion-recognition steps of TC-NER. Another conserved protein ELOF1, or its yeast ortholog Elf1, was recently identified as a core transcription-coupled repair factor. How Rad26 distinguishes between RNA polymerase II (Pol II) stalled at a DNA lesion or other obstacles and what role Elf1 plays in this process remains unknown. Here, we present cryo-EM structures of Pol II-Rad26 complexes stalled at different obstacles that show that Rad26 uses a universal mechanism to recognize a stalled Pol II but interacts more strongly with a lesion-arrested Pol II. A cryo-EM structure of lesion-arrested Pol II-Rad26 bound to Elf1 revealed that Elf1 induces new interactions between Rad26 and Pol II when the complex is stalled at a lesion. Biochemical and genetic data support the importance of the interplay between Elf1 and Rad26 in TC-NER initiation.

## Introduction

Transcription-coupled DNA nucleotide excision repair (TC-NER), a highly conserved sub-pathway of nucleotide excision repair across all three kingdoms of life, is the first line of defense that detects and removes a broad spectrum of transcription-blocking lesions in the transcribed genome (1-7).

As a master TC-NER factor, Cockayne syndrome group B (CSB) protein, or its ortholog Rad26 in *Saccharomyces cerevisiae*, a member of the Swi2/Snf2 family of nucleosome remodeling helicases/ATPases, plays a crucial early role in eukaryotic TC-NER (1-7). During the lesion recognition steps of TC-NER, CSB/Rad26 distinguishes a lesion-arrested Pol II from other types of arrested Pol II and facilitates subsequent recruitment of downstream repair factors, including CSA, UVSSA, and TFIIH (1, 4-6, 8). In addition to its role in TC-NER, CSB/Rad26 also functions as a processivity factor for Pol II arrested in the absence of DNA damage, and regulates a subset of genes crucial for neurological differentiation and development (8-12). Mutations in CSB are linked to Cockayne syndrome, a severe neurodevelopmental disorder characterized by photosensitivity and premature aging (6, 13). Cryo-EM structures have shown that both yeast Rad26 and human CSB bind to the upstream of a stalled Pol II in an evolutionarily conserved manner (8, 14). While these structures provide important insights into the molecular mechanism of eukaryotic TC-NER, the stalled Pol II complexes were prepared in the absence of DNA lesions. This left an important question unanswered: Does CSB/Rad26 recognize the difference between lesion- and non-lesion-arrested Pol II through different initial interactions, or does it use a common initial binding mode, followed by differentiation based on its ability to use its DNA translocase activity to help Pol II bypass only non-lesion barriers?

ELOF1 (human)/Elf1 (yeast *S*.*cerevisiae* ortholog) was recently identified as another essential transcription-coupled repair factor by several groups using genome-scale CRISPR screens against DNA damaging agents (6, 15, 16). Elf1/ELOF1 is a highly conserved transcription elongation factor that binds to a Pol II elongation complex (15, 17-19). Loss of ELOF1 in humans or deletion of Elf1 in yeast leads to UV sensitivity (6, 15). In human cells, ELOF1 is reported to interact with ubiquitin ligase CRL^CSA^ and promote UVSSA binding to lesion-stalled Pol II. Knocking out ELOF1 leads to a decrease in UV-induced Pol II ubiquitylation and UVSSA monoubiquitylation (6, 16). These findings, however, do not explain the evolutionarily conserved role of Elf1/ELOF1 in TC-NER since yeast lacks counterparts of CRL^CSA^ and UVSSA, and Pol II ubiquitylation is not essential in yeast TC-NER(20).

We set out to establish whether Rad26 uses the same mechanism to recognize all stalled Pol IIs, regardless of the nature of the obstacle, and if and how Rad26 and Elf1 function together, mechanistically, in TC-NER. We report four high-resolution cryo-EM structures of Pol II stalled at different obstacles, including a UV DNA lesion, cyclobutene pyrimidine dimer (CPD) lesion. These structures reveal that Rad26 uses a universal approach to recognize a stalled Pol II but interacts more strongly with it in the presence of a lesion. Next, we provide functional evidence supporting a role for Elf1 in promoting recruitment of Rad26 to lesion-arrested Pol II. Finally, we present a cryo-EM structure of both Elf1 and Rad26 bound to a lesion-arrested Pol II. Our structure reveals that the presence of Elf1 leads to new interactions between Rad26 and Pol II absent from all other Rad26-containing structures. Functional studies highlight the importance of these Pol II-Rad26 interfaces in TC-NER. Taken together, these results provide an important mechanistic framework for understanding the functional interplay between two key transcription-coupled repair factors—CSB/Rad26 and ELOF1/Elf1—during TC-NER initiation.

## Results

### Rad26 has a common binding mode for different arrested Pol II complexes

To investigate the structural basis of Rad26 recognition of lesion-arrested and non-lesion arrested Pol II, we solved cryo-EM structures of Pol II-Rad26 complexes stalled either at a CPD DNA lesion, (Pol II(CPD)-Rad26)), or containing a transcription scaffold that mimics a backtracked state after arrest at a non-lesion site (Backtracked Pol II-Rad26) **(Figure 1, Figures S1-4)**. In all structures **(Figures 1B-F)**, as it was the case in our previous structure of a Pol II-Rad26 complex stalled at a non-lesion site (by nucleotide deprivation, **Figure 1B**) (8), Rad26 is bound behind the polymerase near the upstream fork of the transcription bubble and interacts with the protrusion and the wall domain of Rpb2, and the clamp coiled-coil of Rpb1. Similarly, the binding of Rad26 bends the upstream DNA by ∼80° towards the Pol II stalk (Rpb4/7) in all cases. Thus, Rad26 has a common mode of interacting with Pol II regardless of the type of arrest **(Figure 1)**.

**Figure 1.**
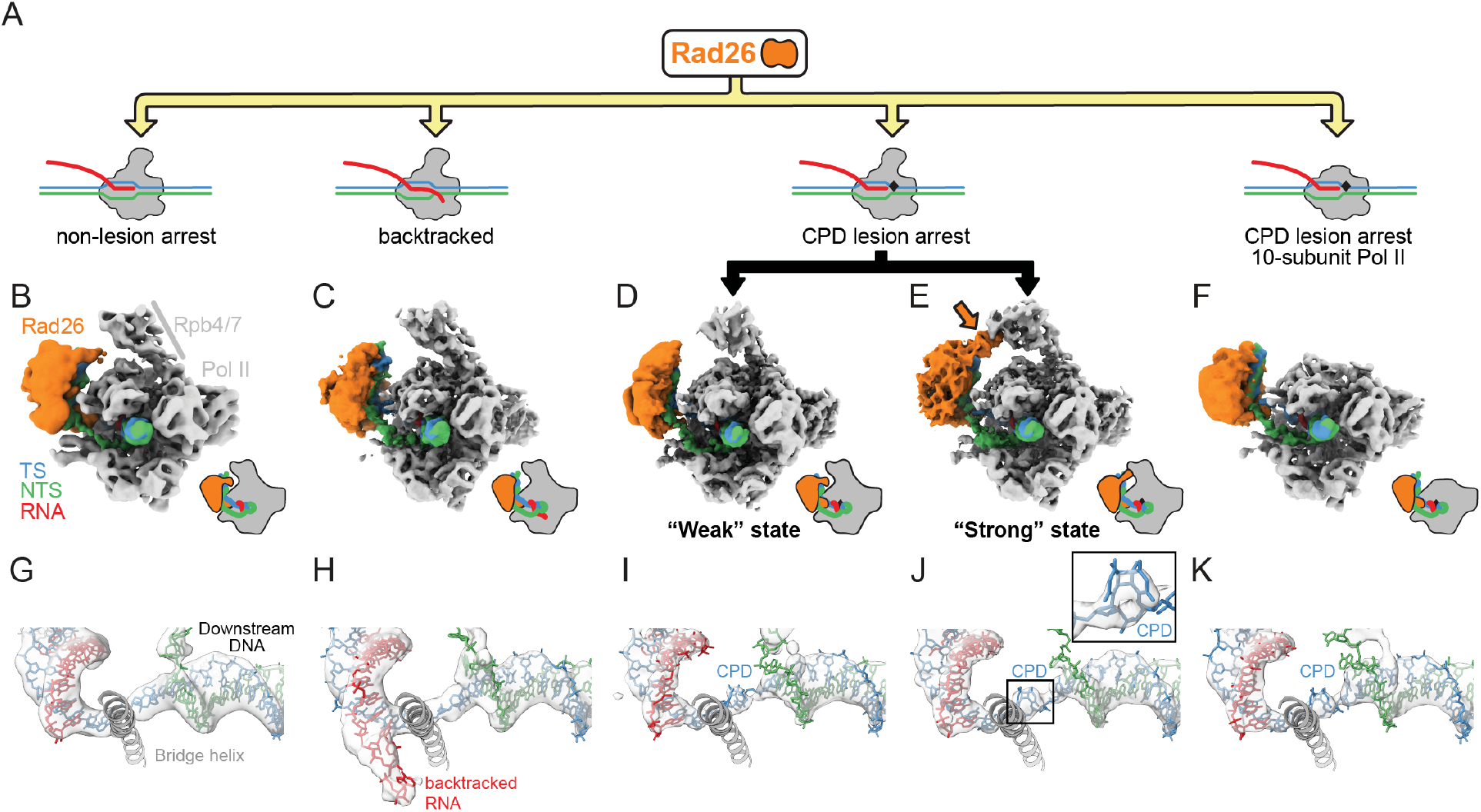
Rad26 interacts in a similar way with paused, CPD-stalled and Backtracked RNA Pol II. **(A)** Cartoon representation of the different complexes analyzed by cryo-EM. The black rhomboid represents the CPD lesion. **(B-F)** Cryo-EM maps of **(B)** Pol II-Rad26 at a non-lesion arrest from our previous work (5.8Å) (8), **(C)** Backtracked Pol II-Rad26 (4.4Å), **(D**,**E)** Pol II(CPD)-Rad26 in states showing either a “Weak” (3.7Å) (D), or “Strong” (3.5Å) (E) interaction between Rad26 and Rpb4/7 (orange arrow), and **(F)** Pol II(CPD)-Rad26 with Pol II lacking Rpb4/7 (4.7Å). The maps were filtered according to the local resolution and were segmented and colored to highlight the different components, as indicated in (B). Cartoon representations of each structure, in the same orientation, are shown next to the maps. **(G-K)** Cryo-EM densities corresponding to the DNA/RNA scaffolds in the vicinity of the active site of Pol II segmented from the maps shown in B-F. The active site Bridge helix was included as a reference point. A close-up of the cryo-EM density corresponding to the CPD lesion is shown in (J). The color scheme used throughout the paper is as follows: Pol II: grey; Rad26: orange; non-template strand: green; template strand: blue; RNA: red.

### A strong interaction between Rad26 and Rpb4/7 is present in the lesion-arrested Pol II-Rad26 complex

Although our structures show that Rad26 uses a common binding mode for all arrested Pol II, differences among them point to arrest-specific interactions between Rad26 and Pol II. When we performed three-dimensional classification of the lesion-arrested Pol II-Rad26 complex dataset (Pol II(CPD)-Rad26) **(Figure S1)**, we found two coexisting conformations. The key difference between them is in the interaction between Rad26 and Rpb4/7 in Pol II: in one state, reminiscent of the structures seen with backtracked and non-lesion-arrested Pol II **(Figure 1B,C)**, and which we termed “Weak”, there is weak density between Rad26 and Rpb4/7 **(Figure 1D)**; in the second, or “Strong” state, there is well defined density connecting them **(Figure 1E)**. This Strong state is specific to the lesion-stalled Pol II(CPD)-Rad26 complex and has three main structural features: Rpb4/7 has shifted towards Rad26 (relative to the core of Pol II); Rad26 has moved towards Rpb4/7, with a concomitant higher bending of the upstream DNA; and, as mentioned above, the density connecting Rpb4/7 and Rad26 is stronger **(Figures 1E, and Figure S5A-C)**. This enhanced Rad26-Rpb4/7 interaction and closer proximity of Rpb4/7 to Rad26 do not appear to be general features of all stalled Pol II complexes as we did not observe them in our previous structures of Pol II-Rad26 stalled at a non-lesion site (8) **(Figure S5D)** or in our new Backtracked Pol II-Rad26 complex **(Figure 1B,C, Figure S6A-C)**. In fact, the Rad26-Rpb4/7 interaction is weakest in the Backtracked Pol II-Rad26 state and Rpb4/7 has moved further away from Rad26 **(Figure S6D)**.

The overall architecture of all Rad26-Pol II (arrested) complexes—the binding to and bending of the upstream DNA— is not dependent on the interaction between Rad26 and Rpb4/7: a structure of core Pol II(CPD)-Rad26 with 10-subunit Pol II showed that the DNA was bent to a similar extent in the absence of Rpb4/7 **(Figure 1F, Figure S7)**. The increased flexibility of Rad26 in this structure **(Figure S7E)**, however, suggests that the interaction of Rad26 with Rpb4/7 is involved in stabilizing the former.

### Elf1 induces new interactions between Rad26 and a lesion-arrested Pol II(CPD)

A previous genome-wide multi-omics analysis of the UV-induced DNA damage response identified human ELOF1 as a top interactor with human CSB (21). Inspired by this, we set out to understand the evolutionarily conserved role of Elf1/ELOF1 in TC-NER at a mechanistic level, focusing on Elf1, the *S. cerevisiae* ortholog of human ELOF1 **(Figures S8A, B)**. We hypothesized that Elf1/ELOF1 could be involved in the initiation of TC-NER by modulating the interaction between Pol II and Rad26/CSB.

Yeast Elf1 and human ELOF1 share a highly conserved core domain **(Figure S8A)**. In addition, Elf1 contains an intrinsic-disordered yeast-specific C-terminus **(Figures S8AB)**, which was not observed in a published structure of Pol II-Spt4/5-Elf1 even though full-length protein was used (19). A yeast strain containing a “core” Elf1 (rad16*Δ*Elf1core) was created by introducing an early stop codon at amino 86 (creating a C-terminal truncation, Elf1*ΔC*) to mimic human ELOF1 **(Figure S8A,B)**. This strain with core Elf1 protein (rad16*Δ*Elf1core) behaved similarly in its response to UV damage *in vivo* compared with the yeast strain (rad16*Δ*) with full-length Elf1 protein **(Figure 2A)**.

**Figure 2.**
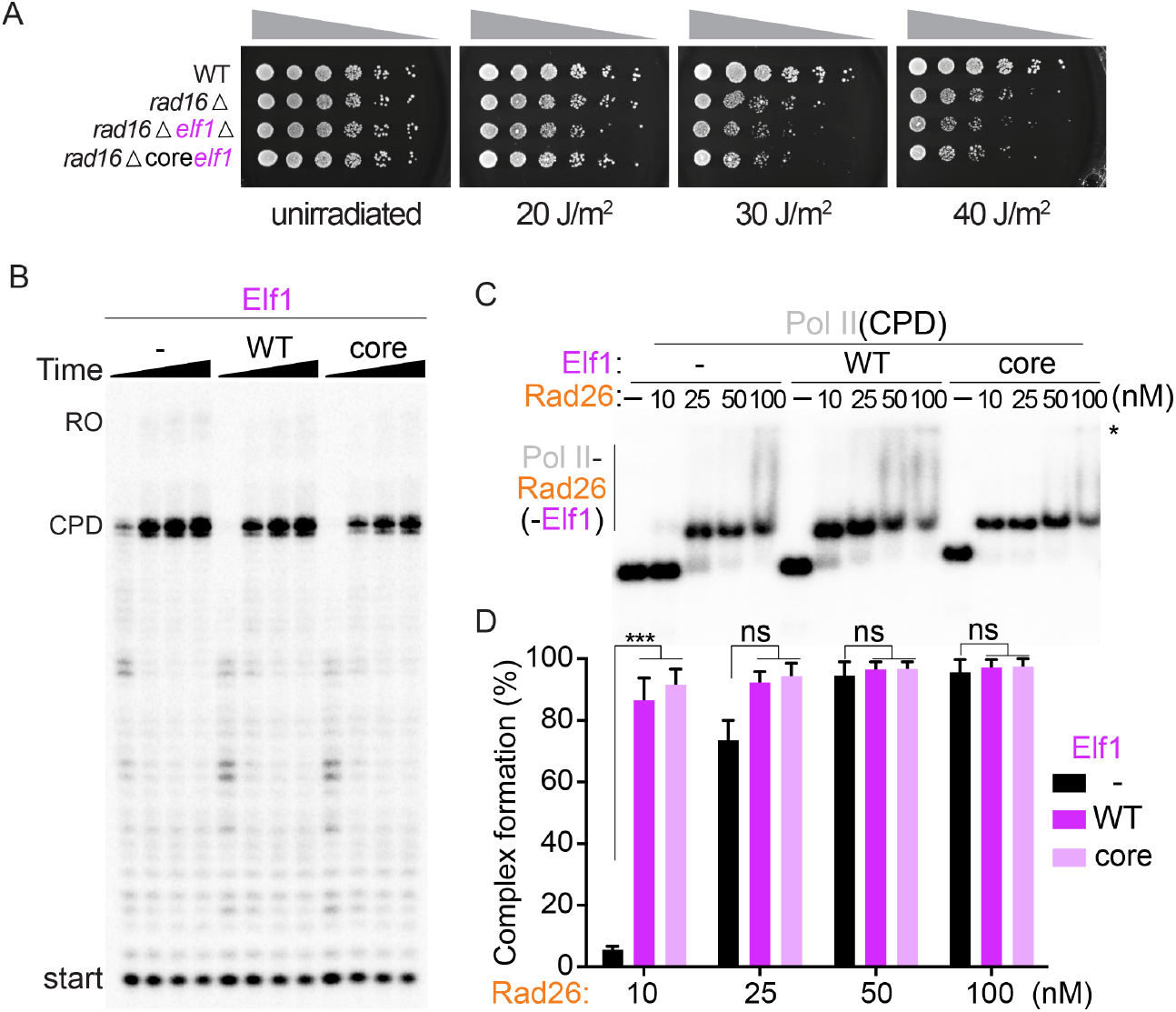
Elf1 enhances interactions between Pol II and Rad26. **(A)** Deletion of Elf1 leads to UV sensitivity and can be rescued by Elf1core. **(B)** Elf1, or Elf1core have no effect on the stalling of Pol II at a CPD lesion, time points were 0.3, 1, 3, and 10 min, respectively. **(C**,**D)** Elf1 enhances the binding of Rad26 to the Pol II complex. The final Elf1 concentration in (B,C) was 1 uM. Data are mean and s.d. (n=3). *** *P* < 0.001, two-tailed Student’s t-test. All assays were performed at least three times independently. The asterisk on the right of the gel in (C) represents the position of the well.

Given what we observed *in vivo*, we tested whether Elf1 (WT) or Elf1*Δ*C proteins behaved in similar ways *in vitro* as well. We first measured their effect on the stalling of Pol II at a CPD lesion in a transcription assay. The stalling pattern of Pol II was the same with either construct **(Figure 2B)**. Next, we tested whether Elf1 (WT) or Elf1*Δ*C affect the Pol II-Rad26 interaction as measured by a gel mobility assay. Intriguingly, we found that both Elf1 (WT) and Elf1ΔC promoted the formation of the lesion-arrested Pol II(CPD)-Rad26 complex (compare 10 nM lanes in **Figure 2C,D**).

To understand the structural basis of Elf1’s role in promoting the interaction between Rad26 and Pol II stalled at a CPD lesion, we solved a cryo-EM structure of a Pol II(CPD)-Rad26-Elf1 complex **(Figure 3A, Figures S9 and S10A-G)**. Elf1 is bound in the downstream channel, next to the lobe domain of Rpb2 and bridging the cleft, as previously reported **(Figure S8C-E)** (18, 19). The presence of Elf1 resulted in a significant improvement in the local resolution of Rad26 to 4Å, from 8Å in the Strong state, our second-best map **(Figure S10A)**. This stabilization effect is likely through a long-range allosteric path (Elf1 ←→ Pol II protrusion domain ←→ Pol II wall domain ←→ Rad26), as there are no direct interactions between Rad26 and Elf1 **(Figure 3A)**. Most strikingly, the Pol II(CPD)-Rad26-Elf1 complex, which we refer to as the “Engaged” state, showed new density at the interface between lobe 2 of Rad26 and the wall domain of Rpb2, corresponding to interfaces absent from any of the other 5 Rad26-Pol II complex structures we have solved to date **(Figure 4, Figure S10H,I)**. The flap-loop of Rpb2, which was disordered in the other structures, is folded, and interacts directly with a short loop-helix-loop region in the Rad26 lobe 2 (631-644aa, which was also disordered in previous structure 5VVR) **(Figure 4)**. This newly folded short loop-helix-loop region is next to the conserved HD-2-1 motif of Rad26, which inserts in the upstream fork of the DNA transcription bubble. The interaction between Rad26 and Rpb4/7 seen in the Strong state is also preserved in this structure **(Figure 3, 4)**. Comparing the bending of the upstream DNA among the Weak, Strong, and Engaged states highlights how Rad26 (and the DNA to which it is bound) shifts towards Pol II as the interactions between them become stabilized **(Figure 3B,C)**. This is consistent with the idea that these structures represent steps in the commitment to fully assemble a lesion-stalled Pol II-Rad26-Elf1 complex for TC-NER.

**Figure 3.**
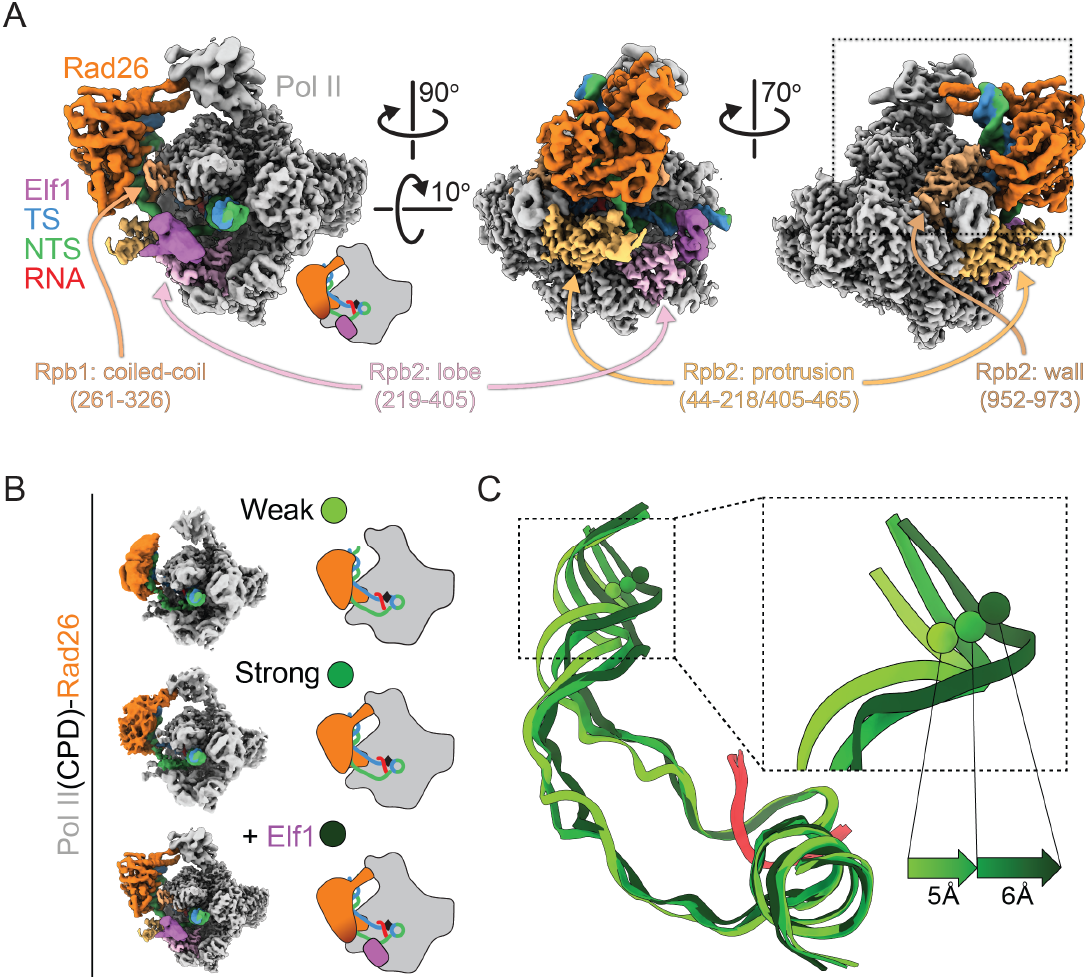
CryoEM structure of Pol II(CPD)-Rad26-Elf1 complex. (A)3.1Å cryo-EM map of the Pol II(CPD)-Rad26-Elf1 complex, with Elf1 colored in light purple. The regions of Pol II that interact with Elf1 or Rad26 are colored in shades of purple or orange, respectively. **(B**,**C)** The increase in the bend angle of the upstream DNA mirrors the stabilization of Rad26 in the cryo-EM maps. **(B)** Cartoon representations of the three structures being compared. **(C)** The DNA/RNA scaffolds were superimposed using the downstream DNA and color-coded as shown on the left. The distances of upstream DNA shift between these states are shown.

**Figure 4.**
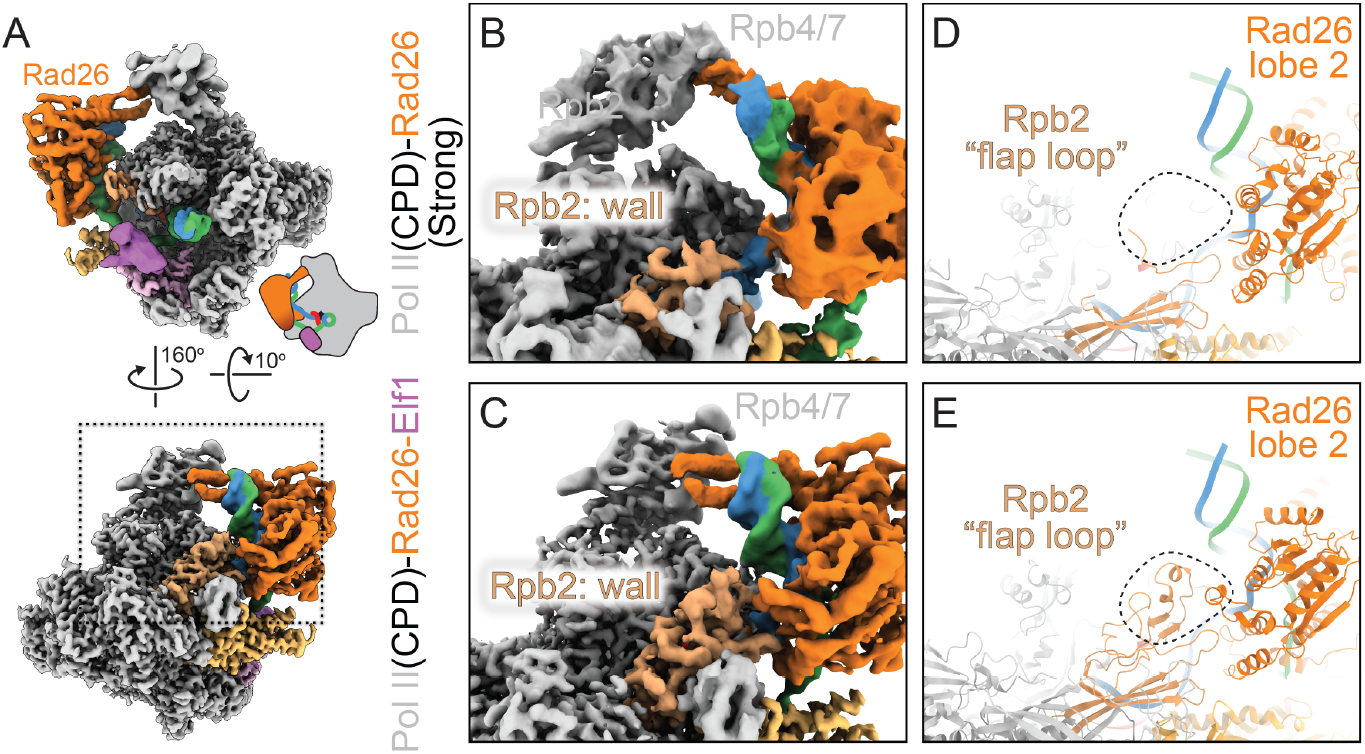
New interaction interfaces between Pol II and Rad26 observed in the Pol II(CPD)-Rad26-Elf1 complex. Overview of Pol II(CPD)-Rad26-Elf1 complex. **(B-E)** Zoomed in views of the area highlighted by the square in **(A)** for Pol II(CPD)-Rad26 without **(B, D)** or with **(C**,**E)** Elf1 bound. **(B**,**C)**: Cryo-EM densities. **(D**,**E)**: Corresponding models. The dashed square highlights the region where new density is present in the Pol II(CPD)-Rad26-Elf1 structure **(E)**. The structural elements that become ordered in both Pol II and Rad26 are shown in (E).

### Pol II-Rad26 interactions stabilized by Elf1 are required for their functional coupling

The resolution of the Pol II(CPD)-Rad26-Elf1 complex allowed us to identify key residues involved in the Elf1-induced Pol II-Rad26 protein interfaces revealed by this structure. One interface is located between a short loop-helix-loop motif of Rad26 lobe 2 (631-644aa, next to HD-2-1 motif) and Pol II Rpb2 wall/flap loop domain (Interface A). The other interface (Interface B) is between Rad26 (475-490aa, connecting conserved motifs IIa (switch) and III in lobe 1) and Rpb1 Clamp coiled-coil domain (Interface B). To further understand the functional significance of these interfaces, we mutated several conserved residues expected to disrupt them: Rad26(R635D/K638D/R639D) (“RKR/DDD”) and Rad26(L483A/K486A/K487A) (“LKK/AAA”). Rad26-RKR/DDD should disrupt the interface between Rad26 and Pol II Rpb2 wall/flap loop domain (Interface A) **(Figures 3A and 4)**, while Rad26-LKK/AAA should disrupt the interface between Rad26 and Pol II Rpb1 Clamp coiled-coil domain (Interface B) **(Figure 3A)**.

We previously showed that Rad26 improves transcription-coupled lesion recognition fidelity and rescues Pol II from non-lesion arrests (8, 11). This relies on coupling Rad26’s ATP-dependent DNA translocase activity with Pol II’s forward translocation to promote Pol II bypass of non-lesion induced arrests. To test whether the Rad26-Pol II interfaces we identified in the Pol II(CPD)-Rad26-Elf1structure are necessary for this function, we purified Rad26-LKK/AAA (Interface B mutant) and Rad26-RKR/DDD (Interface A mutant) and tested their ability to promote Pol II bypass of a pausing sequence. As shown in **Figure 5A**, both Rad26 mutants were significantly impaired in this assay. Importantly, this effect is not due to a reduction in Rad26’s binding to Pol II: both Rad26-RKR/DDD and Rad26-LKK/AAA bind to Pol II with affinities equivalent to that of Rad26(WT) **(Figure 5B). Figure 5B** also shows that the concentrations of Rad26 proteins used in the transcription bypass assay in **Figure 5A** (200nM) were saturating, ruling out the possibility that the effects seen were due to reduced binding of certain Rad26 mutants to Pol II. Another possible explanation for the differences observed in **Figure 5A** was an effect of the mutants on the ATP-dependent DNA translocase activity of Rad26. To test this, we generated and purified the RKR/DDD and LKK/AAA mutants in the context of a constitute-active form of Rad26 (a deletion of the N-terminal auto-inhibitory motif) and characterized their translocase activities using a restriction enzyme accessibility assay with nucleosomes as substrates. All three constructs behaved similarly in this assay **(Figure 5C)**, showing that the effects of the mutations on Rad26’s ability to help Pol II bypass a pausing sequence was not due to different translocation activities. Taken together, these results support a functional role for the Rad26-Pol II interfaces (Interfaces A and B) identified here in coupling Rad26 and Pol II activities for discriminating between lesions and non-lesions.

**Figure 5.**
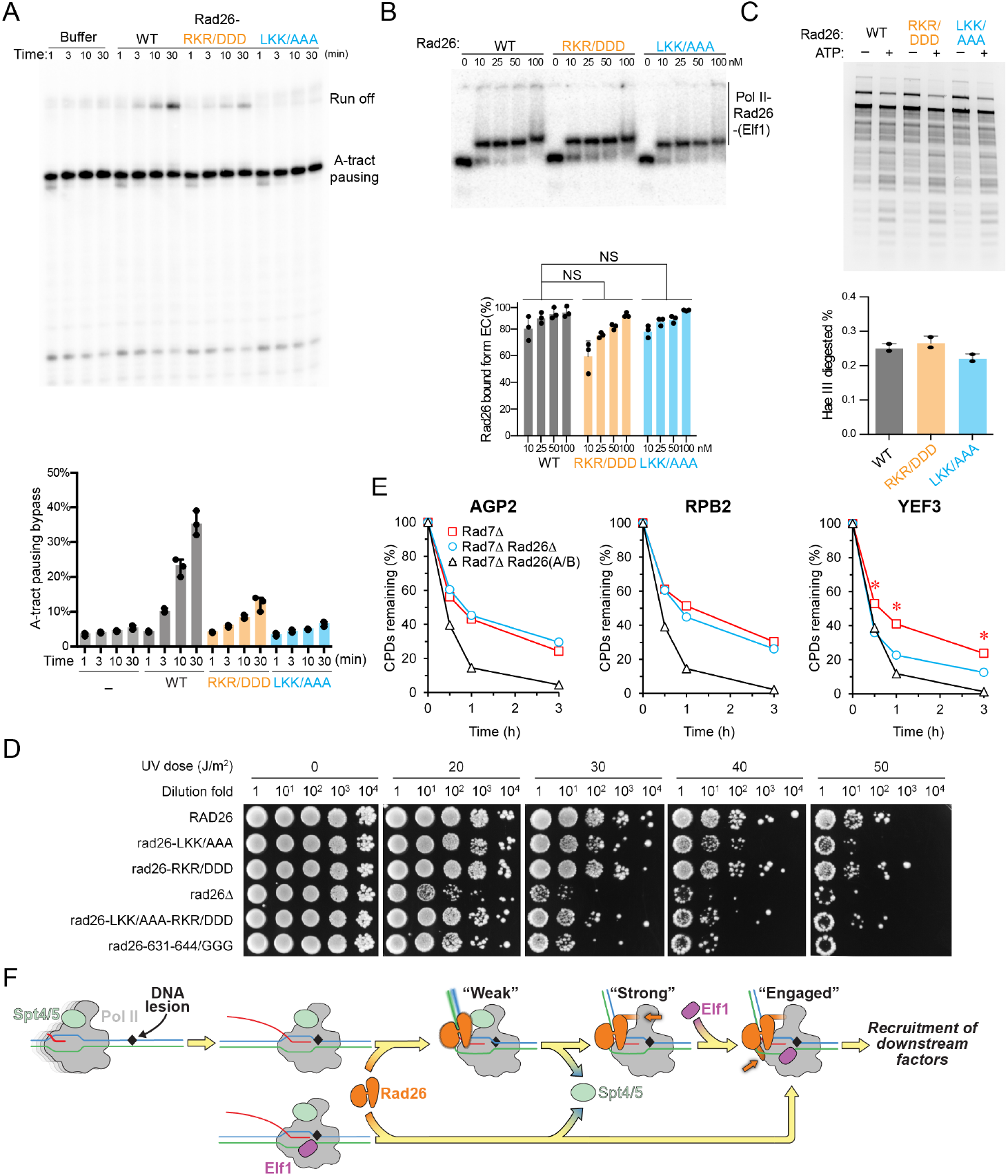
Functional significance of Pol II-Rad26 interface in TC-NER. **(A)** Rad26 mutants (Rad26-LKK/AAA and Rad26-RKR/DDD) are impaired in promoting Pol II transcription bypass of a pausing sequence. **(B)** Rad26-LKK/AAA and Rad26-RKR/DDD do not have impaired binding to Pol II. **(C)** Rad26-LKK/AAA and Rad26-RKR/DDD do not have impaired ATP-dependent DNA translocase activity. All assays in A-C were performed at least three times independently. **(D)** Effects of Rad26 mutations on UV sensitivity. Images are from plates spotted with yeast cells of indicated Rad26 mutations following irradiation with the indicated UV doses. **(E)** Plots showing the effect of *rad26-LKK/AAA-RKR/DDD* on TC-NER. The means of percent CPDs remaining at all CPD sites 50 nucleotides downstream of the major transcription start site (TSS) in the template strand of *AGP2, RPB2* and *YEF3* gens in the indicated cells are plotted. Asterisks (*) indicates that the percent CPDs remaining in the *rad7⊿ rad26-LKK/AAA-RKR/DDD* cells is significantly different from that in the *rad7⊿ rad26⊿* cells at the corresponding repair time points (p values < 10^−10^, Student’s t-test). **(F)** Stepwise model for lesion recognition and reconfiguration during TC-NER initiation in yeast. The color scheme used in Figure 5F is as follows: Elf1: magenta, Spt4/5: lime. Other colors are the same as in Figure 1.

### Mutations of Pol II-Rad26 interfaces abolish TC-NER activity *in vivo*

To test the in vivo role of these interfaces in TC-NER, we first measured the UV sensitivity of the mutants we had designed to disrupt them. The highly efficient Global Genome (GG)-NER in yeast cells would mask the effects of Rad26 on UV sensitivity and TC-NER in cells (22). We therefore used GG-NER-defective rad7*⊿* cells for our assays. As shown in **Figure 5D**, all strains with mutations in Interfaces A and/or B had increased UV sensitivity compared with Rad26(WT), indicating that these mutations render Rad26 TC-NER-defective. Strikingly, we found that three mutations (among all the strains we tested) are most deleterious, increasing UV sensitivity to a level approaching that seen in a rad26 deletion strain: rad26-LKK/AAA; rad26-LKK/AAA-RKR/DDD; or the strain replacing the loop-helix-loop of Rad26 lobe 2 at Interface A with GGG (rad26-631-644/GGG) **(Figure 5D)**.

To further characterize the role of the Rad26-Pol II interactions identified here in TC-NER, we used a well-established TC-NER assay (23) that can quantitively measure the kinetics of TC-NER at different genomic loci at base resolution. We analyzed the effect of Rad26 interface mutations on TCR in three representative genomic loci (*AGP2, RPB2* and *YEF3*), which are transcribed at low, moderate and high rates, respectively (22, 24). In rad7*⊿* cells expressing wild type Rad26, rapid repair of CPDs can be seen immediately downstream of the major transcription start site (TSS) in the template strand (TS) of these genes **(Figure 5E, black curve, and Figure S11)**, indicating rapid TC-NER. TC-NER was slow in rad7*⊿* rad26*⊿* cells **(Figure 5E and Figure S11)**, especially in the region over 50 nucleotides downstream of the transcription start sites (TSS) **(Figure S11)**, where the RNA Pol II switches to transcription elongation mode and TC-NER is repressed by Spt4/5 in the absence of Rad26 (22). Rad7*Δ* cells expressing a Rad26 mutated on both interfaces (rad7*Δ* rad26-LKK/AAA-RKR/DDD) and rad7*Δ* cells lacking Rad26 (rad7*Δ* rad26*Δ*) showed similar rate of TC-NER (slow) in the AGP2 and RPB2 genes **(Figure 5E, compare cyan and red curves, and Figure S11)**, indicating that the Rad26 double interface mutant (rad26-LKK/AAA-RKR/DDD) has no Rad26-dependent TC-NER activity in these genes. Interestingly, the TC-NER rate in the *YEF3* gene was even slower, in a significant way, in rad7*⊿* rad26-LKK/AAA-RKR/DDD cells than in rad7*⊿* rad26*⊿* cells **(Figure 5E, compare red and cyan curves, and Figure S11)**. This suggests that the presence of the Rad26 double interface mutant (rad26-LKK/AAA-RKR/DDD) has a dominant negative effect on TC-NER in the rapidly transcribed gene. Taken together, our *in vitro* and *in vivo* functional data support an important role for the Rad26-Pol II interactions we identified in the Pol II(CPD)-Rad26-Elf1 complexes in TC-NER as well as in coupling the activities of Rad26 and Pol II to allow for the control of lesion recognition/discrimination and the transcriptional bypass of non-lesion arrests.

## Discussion

Here, we report several cryo-EM structures of Pol II-Rad26 complexes arrested at lesion and non-lesion obstacles, revealing three different states: (1) an initial, common binding mode between Rad26 and any arrested Pol II, the “Weak” state, characterized by binding to and bending of the upstream DNA; (2) a lesion-specific “Strong” state with a well-defined interaction between Rad26 and Rpb4/7 in Pol II; and (3) an “Engaged” state, promoted by Elf1, where new interfaces form between Rad26 and Pol II.

### The Weak state represents the initial recruitment of Rad26 to arrested Pol II

A common binding mode of Rad26 to different forms of arrested Pol II is consistent with the dual roles of Rad26 in repair (improving lesion recognition fidelity) and transcription elongation. The Weak state represents the initial recruitment of Rad26 to an arrested Pol II, regardless of the nature of that arrest. At this step, Rad26 is recruited to the upstream DNA fork of arrested Pol II, where it inserts its conserved W752 at the upstream edge of the transcription bubble. Rad26 then uses its ATP-dependent translocase activity to track along the template strand in a 3’-5’ direction, moving towards Pol II. If Pol II is arrested at a non-lesion or small lesion transcription barrier, Rad26’s translocation can promote Pol II forward translocation and eventually obstacle bypass. If, instead, Pol II is arrested at a bulky lesion, bypass cannot occur and the interfaces between Pol II and Rad26 strengthen to form the Strong, and eventually Engaged states. The ultimate consequence of these structural changes is the recruitment of downstream repair factors.

Recently, a lower resolution cryo-EM structure of *Komagataella pastoris* Pol II-Rad26-Elf1 elongation complex on a nucleosome template was published (PDB 8HE5) (25). Although the authors found that Rad26 binds upstream of Pol II, the conformation of *K. pastoris* Pol Rad26 is different from that in the previously reported yeast Pol II-Rad26 and human Pol II-CSB complexes, as well as in the structures presented in this manuscript. The interactions between Pol II and Rad26 are less extensive and may therefore represent an early intermediate preceding the Weak state described here.

### A lesion arrest induces a strong interaction between Rad26 and Pol II stalk

Our structural analysis revealed that a well-defined interaction is formed between Rad26 and the Pol II stalk (Rpb4/7) when Pol II is arrested at a CPD lesion (the Strong state) **(Figure 2E)**. Rpb4/7 plays an important role in controlling the conformational dynamics of the Pol II clamp and is a hub for interactions with several transcription factors, including Spt4/5 and Spt6. As a result, Rpb4/7 plays important roles in several molecular and cellular processes, including transcription and DNA repair (18, 26). A previous genetic study in *S. cerevisiae* showed that Rpb4/7 promotes Rad26-dependent TC-NER while suppressing Rad26-independent TC-NER (27). On the other hand, Spt4/5, an elongation factor that binds both Rpb4/7 and the protrusion domain of Pol II (18, 19, 28), functions as an inhibitor of TC-NER (29). We propose that the steric exclusion of Spt4/5 by the lesion-induced strong interaction between Rad26 and Rpb4/7 is a major step in committing a lesion-stalled Pol II to TC-NER. Furthermore, a previous computational study suggested that tethering Rad26-NTD with Pol II Rpb4/7 is important for Pol II-Rad26 complex assembly and plays a key role in anchoring Rad26 to Pol II and establishing a productive orientation with respect to the transcription bubble (30).

### Binding of Elf1 promotes the formation of Rad26-Pol II interactions critical for TC-NER

Addition of Elf1 increased the affinity of Rad26 for Pol II *in vitro* and resulted in new interactions between Rad26 and Pol II seen in the cryo-EM structure of the Rad26-Pol II(CPD)-Elf1 complex that were absent from all other CPD-stalled structures we have solved to date. Increased affinity is also consistent with the significant improvement in the density for Rad26 in the structure. Binding of Elf1 to Pol II encircles the downstream DNA within the cleft and reduces the conformational dynamics of the Pol II clamp. Given the absence of direct interactions between Elf1 and Rad26, we hypothesize that Elf1 exerts its effect on the Rad26-Pol II interaction through long range allostery. This idea is supported by predictions from a dynamic network analysis (30).

The Pol II-Rad26 interfaces induced by Elf1 are critical for TC-NER; disrupting these interfaces completely abolished TC-NER *in vivo* **(Figure 5E)**. Interestingly, at the highly transcribed *YEF3* gene, we found that a Rad26-Pol II interface mutant led to even slower TC-NER than the cells with deletion Rad26 **(Figure 5E)**, suggesting that the Rad26-Pol II interface mutant protein not only affects Rad26-depedent TC-NER, but also sterically blocks Rad26-indepedent TC-NER at certain gene loci. Taken together, our data suggest that the Pol II(CPD)-Rad26-Elf1 complex represents the fully assembled “engaged” complex in TC-NER.

### Conservation and differences between yeast and human TC-NER initiation

Core TC-NER factors are highly functionally and structurally conserved between yeast and human. Indeed, previous studies revealed striking structural similarity between yeast Pol II-Rad26 and human Pol II-CSB complexes as well as their mechanisms (8, 14). Our study sheds important mechanistic insights into the conserved roles of TC-NER factors Rad26/CSB and Elf1/ELOF1 for TC-NER initiation. Considering the data presented here, along with previous work, we propose a stepwise model for the initiation of TC-NER **(Figure 5F)**. Initial interaction of Rad26 with an arrested Pol II results in bending of the upstream DNA and displacement of Spt4/5 by blocking its interaction with the Rpb4/7 stalk. Rad26’s remodeler-like DNA translocation biases Pol II forward, promoting the bypass of non-lesion obstacles or small lesions (8, 11). In the case of lesion-arrested Pol II, the interaction between Rad26 and Rpb4/7 is strengthened. Binding of Elf1 induces new interactions between Rad26 and Pol II, setting the stage for the recruitment of downstream factors, such as TFIIH and XPA, for TC-NER (14, 25, 31, 32) These initial steps are likely conserved between yeast and human, though additional factors and layers in human cells are involved in regulating TC-NER. For example, CSA and UVSSA, for which there are no yeast counterparts, are important for regulating Pol II ubiquitinylation and in turn recruitment of TFIIH (33).

## Acknowledgements

This work was supported by grants from the National Institutes of Health (R01 GM102362 and R01 GM092895 to D.W; R01 GM092895 and R35 GM145296 to A.E.L.; R01 GM111458 to N.H.; and NIH grant R03 ES033789 to S.L.). The work was also supported by NSF grant MCB-2102072 (to S. L.). We also thank the UC San Diego Cryo-EM Facility, and the UC San Diego Physics Computing Facility for IT support. We are grateful to Dr. Daniel Durocher’s lab for providing yeast strains.

## Author contributions

D.W. and A.E.L. conceived the project. J.X. J.O. and J.C. purified proteins. J.X. assembled complexes for cryo-EM samples and biochemistry assays. J.X. performed biochemical assays. R.D.S. and I.L. performed the cryo-EM work. R.D.S., I.L. J.X., J.O. built atomic models. Z.Z. and N.H. created new yeast strains with Elf1 C-terminus truncation and performed genetic experiments. W.G. and S. L. created new yeast strains with Rad26 interface mutations, performed genetic experiments, and measured TC-NER activities of these strains. R.D.S., J.X., I.L. Z.Z., N.H., W.G., S. L., A.E.L., and D.W. performed data analysis. D.W. and A.E.L. supervised the different aspects of the work. R.D.S., J.X., I.L., J.C., S. L., A.E.L, and D.W. made the figures and wrote the manuscript, with input from all authors.

## Data and materials availability

Cryo-EM maps and corresponding models have been deposited in the EM Data Bank and Protein Data Bank, respectively. Accession codes can be found in Table S1.

## Materials and Methods

### Materials and Methods

#### Protein expression and purification

Expression and purification of Rad26 were performed essentially as previously described (3). Briefly, recombinant Rad26 protein was expressed in *Escherichia coli* strain Rosetta 2(DE3) (Novagen) and purified by Ni-NTA agarose (Qiagen), Hi-Trap Heparin HP (GE Healthcare), and Superdex 200 10/300 GL columns (GE Healthcare). Rad26 mutants were purified in the same manner as wild-type proteins. Expression and purification of yeast TFIIS were performed as described (4). Expression and purification of yeast Elf1 and yeast Elf1core were performed essentially as previously described (5, 6). Briefly, GST-tagged Elf1 protein was expressed in *Escherichia coli* strain Rosetta 2(DE3) (Novagen) and purified by Glutathione Sepharose 4 Fast Flow resin (GE Healthcare), and Superdex 200 10/300 GL column (GE Healthcare). Elf1core was purified in the same manner as wild-type protein. Recombinant Spt4/5 was a gift from Dr. Jianhua Fu and expressed and purified as described (7).

*Saccharomyces cerevisiae* 10-subunit Pol II was purified essentially as previously described (8). Briefly, Pol II (with a protein A tag in the Rpb3 subunit) was purified by an IgG affinity column (GE Healthcare), followed by Hi-Trap Heparin (GE Healthcare) and Mono Q anion exchange chromatography columns (GE Healthcare). Pol II was purified by incubating 10-subunit Pol II with 3-fold of Rpb4/7 followed by gel filtration. His6-tagged Rpb4/7 heterodimer was purified from E. coli by Ni-affinity chromatography followed by gel filtration as previously described (9).

#### In vitro transcription assay

Pol II elongation complexes were assembled essentially as previously described with a labeled RNA primer (3). For transcription assay to test the effect of Elf1 or Rad26, purified Elf1 (300 nM or 1 μM) or Rad26 proteins (100 or 200 nM) were also included in transcription assays. *In vitro* transcription was started by adding rNTPs mixture to a final concentration of 1 mM each and quenched at different time points. The reacted samples were boiled for 10 min at 95°C in formamide loading buffer, and the RNA transcripts were separated by denaturing PAGE (6M urea). The gel was visualized by phosphorimaging and quantified using Image Lab software (Bio-Rad).

#### Preparation of Pol II-Rad26 and Pol II(CPD)-Rad26-Elf1 complexes for electron microscopy

Template and non-template DNA oligonucleotides were obtained from IDT and further purified by PAGE. PAGE-purified RNA oligonucleotides were purchased from Dharmacon. HPLC-purified CPD lesion-containing template was purchased from TriLink. The RNA, template DNA (non-damaged or CPD lesion containing) and non-template DNA were annealed to form the scaffold as previously described (3).

To form the CPD-arrested Pol II complex, Pol II and three-fold excess of scaffold were mixed and further purified by gel filtration in 50mM HEPES, pH7.4, 5mM DTT, 5mM MgCl_2_, and 40mM KCl. To form the backtracked Pol II complex, Pol II and the scaffold were incubated in 50mM HEPES, pH7.4, 5mM DTT, 5mM MgCl_2_, and 40mM KCl. To form the backtracked Pol II-Rad26 complex, Rad26 was added to backtracked Pol II complex and incubated for 30 minutes. The final buffer was composed of 50mM HEPES, pH 7.4, 5mM DTT, 5mM MgCl_2_, 40mM KCl, 200mM NaCl. For the Pol II(CPD)-Rad26 complex, a final concentration of 0.02% Glutaraldehyde was added after adding Rad26 and incubated for another 30 minutes. The crosslink reaction was terminated by adding 1M Tris-HCl, pH 8.0 to a final concentration of 100mM. The final concentrations of the different components were 1*μ*M Pol II, 2*μ*M Rad26, 1.1*μ*M scaffold. To form the Pol II(CPD)-Rad26-Elf1 complexes, 4-fold excess of Elf1 was incubated with Pol II(CPD)-Rad26 complex. The final concentrations of the different components were 1*μ*M Pol II, 2*μ*M Rad26, 1.1*μ*M scaffold, and 5*μ*M Elf1.

The sequences used for elongation complex preparation are as follows: non-template DNA, 5′-CTAGTTGATCTCATATTTCATTCCTACTCAGGAGAAGGAG-CAGAGCG-3′; template DNA, 5′-CGCTCTGCTCCTTCTCCCATCCTCTCGATGGCTATGAGATCAACTAG-3′; CPD lesion-containing template DNA, 5′-CGCTCTGCTC CTTCTCCXXTCCTCTCGATGGCTATGAGATCAACTAG-3′ (XX = CPD lesion); RNA (for Pol II(CPD)), 5′-AUCGAGAGGA-3′; RNA (for Back-tracked Pol II), 5′-AUCGAGAGGAUGCAGAC-3′.

#### Electron microscopy

An aliquot of 4uL of each sample was applied to glow-discharged Quantifoil holey carbon films R1.2/13 Cu grids. The grids were blotted and plunge-frozen in liquid ethane using a Vitrobot Mark IV (FEI). Data collection was performed using Leginon ^(10)^ on an FEI Talos Arctica operated at 200 kV, equipped with a Gatan K2 summit direct detector. For the Pol II(CPD)-Rad26 sample, 3,358 movies were recorded in counting mode at a dose rate of 11.3 electrons pixel^-1^s^-1^ with a total exposure time of 7.05 s sub-divided into 150 ms frames, for a total of 47 frames. The images were recorded at a nominal magnification of 36,000x resulting in an object-level pixel size of 1.16 Å pixel^-1^. For the Backtracked Pol II-Rad26 sample, 9,167 movies were recorded in counting mode at a dose rate of 6.75 electrons pixel^-1^ s^-1^ with a total exposure time of 11 s sub-divided into 200 ms frames, for a total of 55 frames. The images were recorded at a nominal magnification of 36,000x resulting in an object-level pixel size of 1.16 Å pixel^-1^. For the Pol II(CPD)-Rad26 sample with Pol II lacking Rpb4/7, 955 movies were recorded in super-resolution mode at a dose rate of 5.34 electrons pixel^-1^ s^-1^ with a total exposure time of 13 s sub-divided into 250 ms frames, for a total of 44 frames. The images were recorded at a nominal magnification of 36,000x resulting in an object-level pixel size of 1.16 Å pixel^-1^ (0.58 Å per super-resolution pixel). For the Pol II(CPD)-Rad26-Elf1 sample, two datasets with total of 6,000 movies were recorded in counting mode at a dose rate of 6.9 electrons pixel^-1^ s^-1^ for the first dataset and 7.4 electrons pixel^-1^ s^-1^ for the second dataset with a total exposure time of 10 s sub-divided into 200 ms frames, for a total of 50 frames. The images were recorded at a nominal magnification of 36,000x resulting in an object-level pixel size of 1.16 Å pixel^-1^. See Table S1 for details on cryo-EM data collection, refinement and validation.

#### Image processing

Movie frame alignment was performed using MotionCore2 ^(11)^ using the dose-weighted frame alignment option. CTF estimation was executed on the non-dose-weighted aligned micrographs using GCTF using the local defocus per particle option (12). Particle picking was performed using FindEM ^(13)^ with 2D averages selected from the initial processing serving as templates. Motion correction, CTF estimation and particle picking were performed within the framework of Appion (14). Two-dimensional classification was performed to identify bad Pol II particles. Following 2D classification, an initial 3D classification was performed using a Pol II Elongation Complex model (PDB 1Y77) as reference. The 2D and initial 3D classifications were carried out using particles binned by 4 (4.64 Å pixel^-1^). The detailed processing schemes for each sample are shown in Figures S1, S3, S7, S9. All initial refinements and classifications were done in Relion 3(15). Once the final particles were selected, local and global ctf refinement were performed to further improve the resolution using cryoSPARC(16) The final map was refined in cryoSPARC using non-uniform refinement algorithm(17). The statistics for refinement of all maps are listed in Table S1,2.

#### Model building

For building the models of Pol II (CPD) Conformation 1 and 2, models of Pol II(CPD) complex (PDB accession 6O6C)(18) and Rpb4/7 of Pol II elongation complex model (PDB accession 5VVS)(3) were used as starting models for Pol II core (10 subunits) and Rpb4/7, respectively. The composite reference model of Pol II core and Rpb4/7 and the density maps were used as inputs in RosettaCM(19), in which 10 models were generated. A model with the best Rosetta energy was selected for each density map. Models were manually optimized in coot(20) and then refined using Rosetta Relax to further optimize the position and geometry of the amino acids side chains. The nucleic acids scaffold was manually built in coot. A selected model was refined using PHENIX real space refinement (21) with secondary structure restrains option followed by second round of Rosetta Relax, in which 10 models were generated. A model with the best score function was selected as the final model. The metals were manually added to each model followed by a final run of PHENIX real space refinement. The model of backtracked Pol II complex apo was built using the same steps described above except that the Pol II(CPD) complex Conformation 1 model was used as a starting model.

For building the model of Rad26 for Pol II(CPD)-Rad26, Backtracked Pol II-Rad26 complex, and Pol II(CPD)-Rad26 with Pol II lacking Rpb4/7, the model of Pol II-Rad26 stalled (PDB accession 5VVR) (3) was used as a reference. The Rad26 starting model was rigid body docked into the density map using UCSF Chimera(22). The N-terminal helix of Rad26 was manually adjusted or deleted in coot to best fit the density map. The composite reference model of optimized Rad26 and Pol II(CPD) Conformation 1 (built as described above) was used as a starting model in RosettaCM.

To build the model Pol II(CPD)-Rad26-Elf1, the reference models for Rad26 and Elf1 were selected based on homology detection using hidden Markov model as implemented in HHpred (23). The segmented density of Rad26 and Elf1, and the references models from HHpred were used as inputs to build their models using RosettaCM. Pol II was built using the composite models of Pol II 10 subunit (from PDB: 6O6C) and Rpb4/7 (from PDB: 5VVS) as described above. Nucleic acid scaffolds for all models were built in coot. The same steps described above were performed to improve position and geometry of the amino acids side chains. FSC curves of map-to-model were calculated in Rosetta. The validation statistics for all models are shown in Table S1.

#### Structure analysis

All figures were generated using UCSF ChimeraX(24). The cryo-EM maps were first segmented using Seggar(25) as implemented in UCSF Chimera. The segmented densities were colored in ChimeraX.

To generate the difference map for Strong state minus Weak state of Pol II(CPD)-Rad26, the cryo-EM maps were first low-pass filtered in SPIDER(26) with FQ operation, ‘top-hat’ function preserving frequencies below 0.1 (a resolution of 10Å in our maps). The difference map was generated in ChimeraX with volume operation (vol) as follows: the filtered Strong state was fitted into Weak state map with ‘fitmap’ command, the Strong state was resampled on the grid of Weak state map with ‘vol resample’ command, and the Weak state map was subtracted from the resampled Strong state map with ‘vol subtract’ command. The same steps were followed to generate difference map for Pol II(CPD)-Rad26 (Strong state) minus Backtracked Pol II-Rad26 and Pol II(CPD)-Rad26-Elf1 minus Pol II(CPD)-Rad26.

The consensus refinement and the masks used in multi-body refinement were prepared in Relion 3 using the default options. Multi-body refinement generated 10 structures, which describe flexibility along each eigenvector. To visualize flexibility along eigenvector 1 and 2 for Pol II(CPD)-Rad26 (Strong state), the model of Rpb4/7 was segmented out from Pol II(CPD)-Rad26 Strong Rad26-Rpb4/7 state model and was rigid-body fitted separately into each one of the ten structures from multi-body refinement. The models were fitted using ‘fitmap’ command in ChimeraX. The same steps were followed to visualize the flexibility along eigenvector 1 and 2 for Pol II(CPD) but using the consensus refoment of Pol II(CPD) to generate masks and as an input for multi-body refinement. The segmented model of Rpb4/7 chains from Pol II(CPD) (lacking Rad26) model was used for rigid-body fitting.

To obtain the cross-correlation coefficients between the Rpb4/7 model and the different cryo-EM maps shown in Figure S5, the cryo-EM maps for the Strong and Weak states of Pol II(CPD)-Rad26 were aligned with the ‘fitmap’ function in ChimeraX. Then, the full complex model was aligned to its corresponding cryo-EM density. To calculate cross-correlation coefficients, the model of Rpb4/7 from Pol II(CPD)-Rad26 (Strong state) was fitted into the segmented Rpb4/7 density from Weak and Strong state while disabling the options for allowing any rotations and shifts. The same steps were performed to calculate the cross-correlation for the fitting of Rpb4/7 model for Strong state of Pol II(CPD)-Rad26 into the map of Backtracked Pol II-Rad26 (Figure S6).

#### Yeast strain construction

Yeast strains used in this study are listed in Table S2. Plasmids expressing 6*⊿*FLAG tagged wild type (p6FRAD26) and indicated mutant Rad26 proteins were created using the pRS415 vector (27). Yeast strains expressing 6*×*FLAG tagged wild type and mutant Rad26 proteins were created by transforming the yeast strain CR18 (28) with the plasmids.

To make *elf1-*▫*C* mutant strain, a *URA3* fragment from pRS306 was PCR amplified with Primer 1 and Primer 2 (see below) to replace the part of *ELF1* open reading frame that encodes C-terminal 60 amino acids (with the incorporation of a TAA stop codon immediately after the amino acid 58). The PCR-cassette was transformed into cells using the method described previously (29). The resulting mutant strains were further confirmed by sequencing. Primers are listed below:

Primer 1: TGATGTATATAGTGATTGGTTTGACGCCGTCGAAGAAGTCAATTCTGGCCGTGGATAACCTGATGCGGTATTTTCTCC.

Primer 2: TTAAAATATAAAATATATATGACCTAAGTAAATATGGTTTTTTCTCAGGACCGGACGGCATCAGAGCAGATTGTA

All genotypes of yeast strains are listed in Table S2.

#### Mapping repair of UV induced CPDs

Yeast cells were cultured at 30°C to late log phase (OD600 ≈ 1.0), irradiated with 120 J/m2 of ∼254 nm UV and incubated at 30°C. At different timepoints of the post-UV incubation, aliquots were taken, and the genomic DNA was isolated. The CPDs remaining in the AGP2, RPB2 and YEF3 genes in the isolated genomic DNA were analyzed using the Lesion-Adjoining Fragment Sequencing (LAF-Seq) method (30). Sequencing reads whose 3’ ends adjoin the sites ofCPDs remained in the genomic DNA were aligned to the TS and/or NTS sequences of the AGP2, RPB2 and YEF3 gene fragments. Reads corresponding to CPDs at individual sites along the gene fragments were counted after subtraction of the background counts (in the unirradiated samples) by using codes in R. To better illustrate the CPD induction and repair profiles, pseudo images whose band intensities correspond to the counts of aligned sequencing reads were generated using codes in R.

#### UV survival assay

Yeast cells were grown at 30°C to optical density (OD) of 3 at 600 nm and diluted to OD 0.6 in YPD medium. Cells were plotted on YPD plate with 5-fold (Figure 2) or 10-fold (Figure 5) serial dilutions. Once dried, the plates were UV irradiated with UV crosslinker (FisherBiotech FB-UVXL-1000) in dark room and wrapped in foil after irradiation. Plates were incubated in the dark for 2-5 days at 30°C before imaging.

---

**Figure S1.**
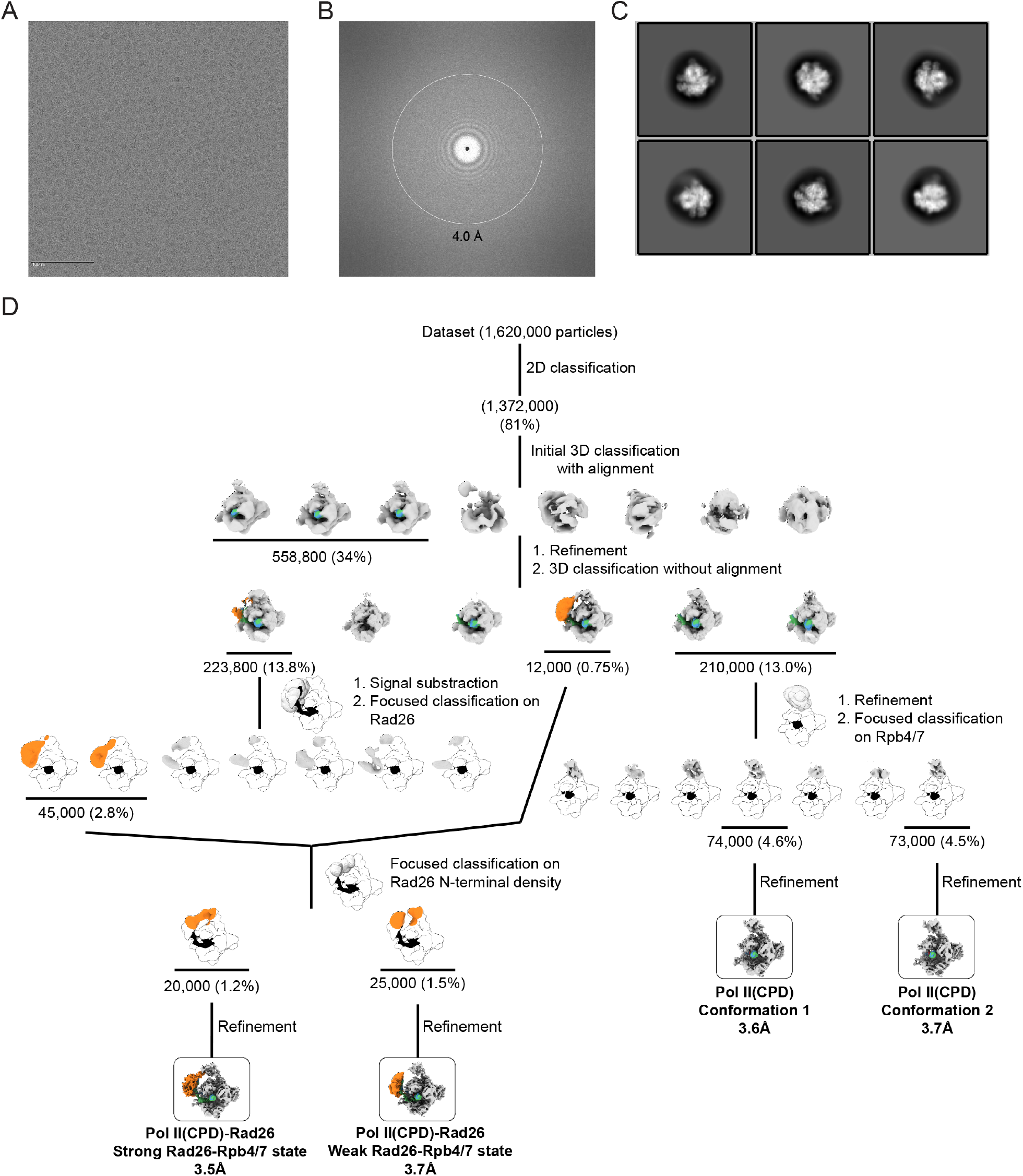
Cryo-EM structure determination of the Pol II(CPD)-Rad26 and Pol II(CPD) complexes. **(A-C)** Representative micrograph **(A)**, power spectrum **(B)**, and representative 2D class averages **(C)** of Pol II(CPD)-Rad26 complexes. **(D)** Schematic of the strategy used to sort out the dataset into Pol II(CPD)-Rad26 Strong and Weak Rad26-Rpb4/7 states, and Pol II(CPD) conformations 1 and 2. Focused 3D classification was performed without alignment unless otherwise noted. The number of particles contributing to each selected structure is indicated. The percentages shown are related to the total number of particles picked from the micrographs. The indicated resolution corresponds to the 0.143 Fourier shell correlation (FSC) based on gold-standard FSC curves (see Figure S2).

**Figure S2.**
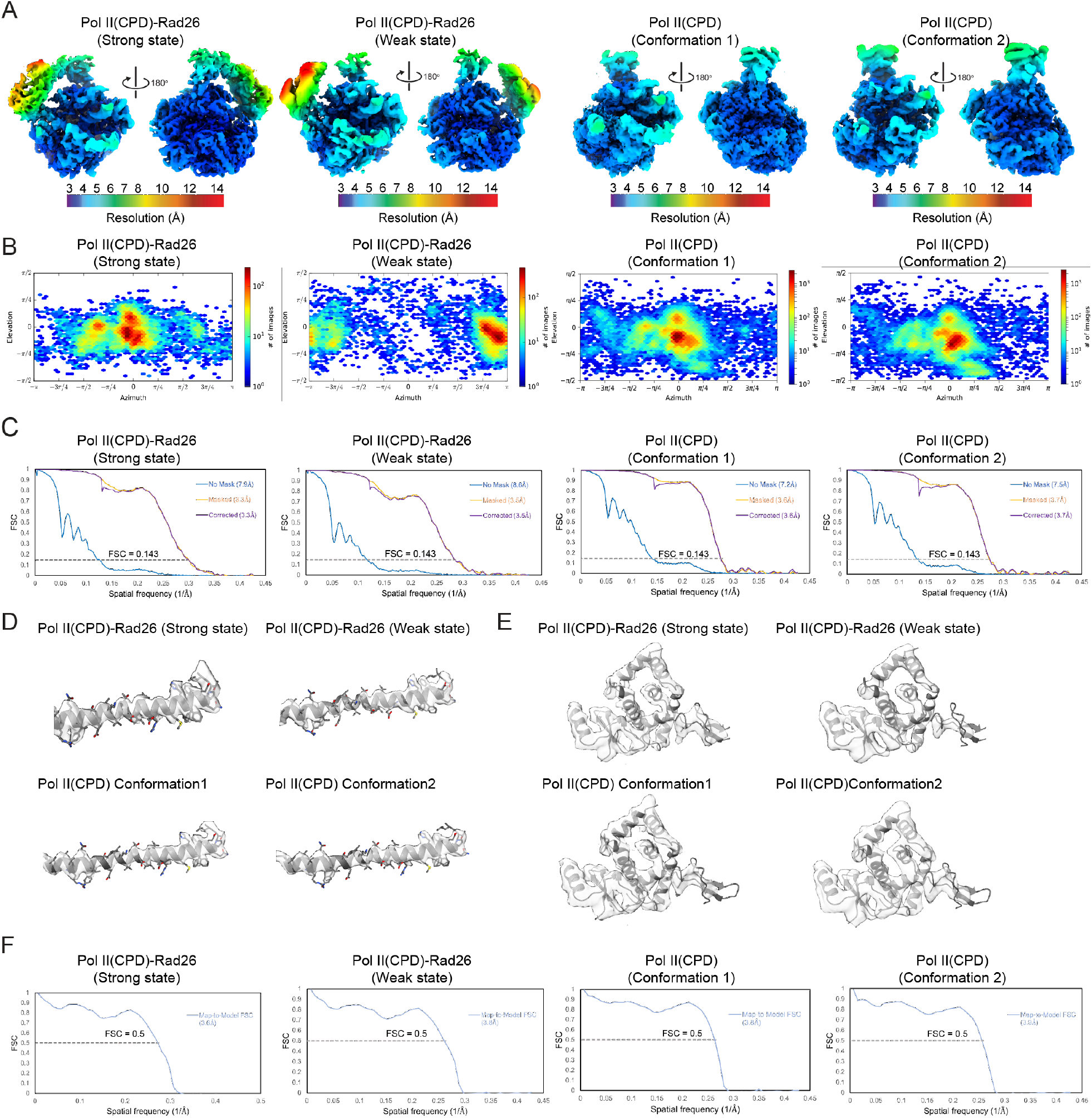
Analysis of the Pol II(CPD)-Rad26 and Pol II(CPD) cryo-EM maps. **(A)** Front and back views of locally filtered maps, colored by local resolution, of Pol II(CPD)-Rad26 Strong and Weak states, and Pol II(CPD) conformations 1 and 2. **(B, C)** Euler angle distribution of particle images **(B)** and FSC plots **(C)** for the maps shown in (A). **(D-E)** Close-ups of the cryo-EM densities corresponding to the Rpb1 Bridge helix **(D)**, and the Rpb2/Rpb9 ‘Jaw’ of Pol II **(E)** for the indicated structures with the models fitted in. **(F)** FSC curves for map-to-model fits for the maps shown in (A). The 0.5 FSC line is shown.

**Figure S3.**
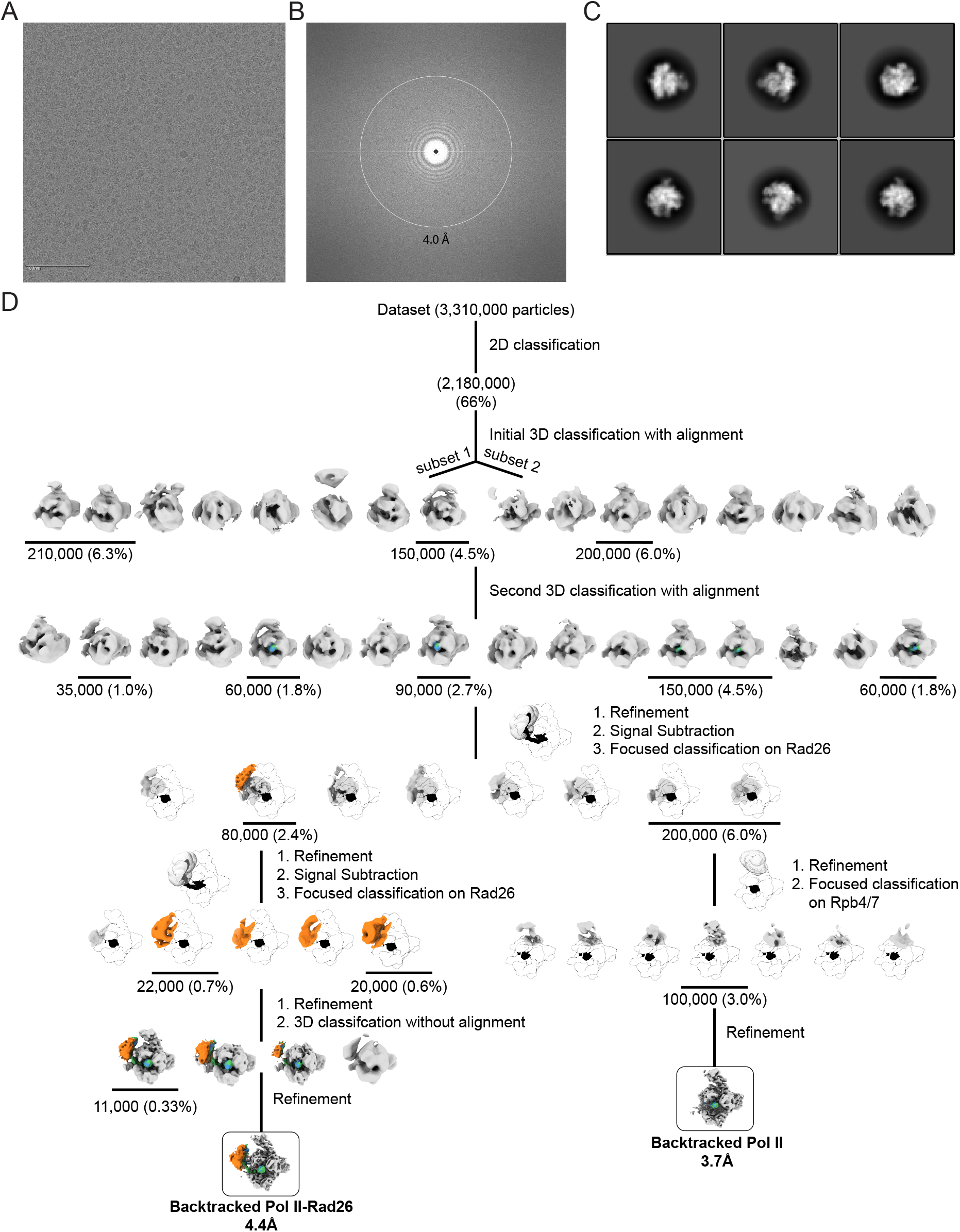
Cryo-EM structure determination of the Backtracked Pol II-Rad26 and Backtracked Pol II complexes. **(A-C)** Representative micrograph **(A)**, Power spectrum **(B)**, and representative 2D class averages **(C)** of Backtracked Pol II-Rad26 complexes. **(D)** Schematic representation of the strategy used to sort out the dataset into Backtracked Pol II-Rad26 and Backtracked Pol II. Focused 3D classification was performed without alignment unless otherwise noted. The number of particles contributing to each selected structure is indicated. The percentages shown are related to the total number of particles picked from micrographs. The indicated resolution corresponds to the 0.143 Fourier shell correlation (FSC) based on gold-standard FSC curves (see Figure S4).

**Figure S4.**
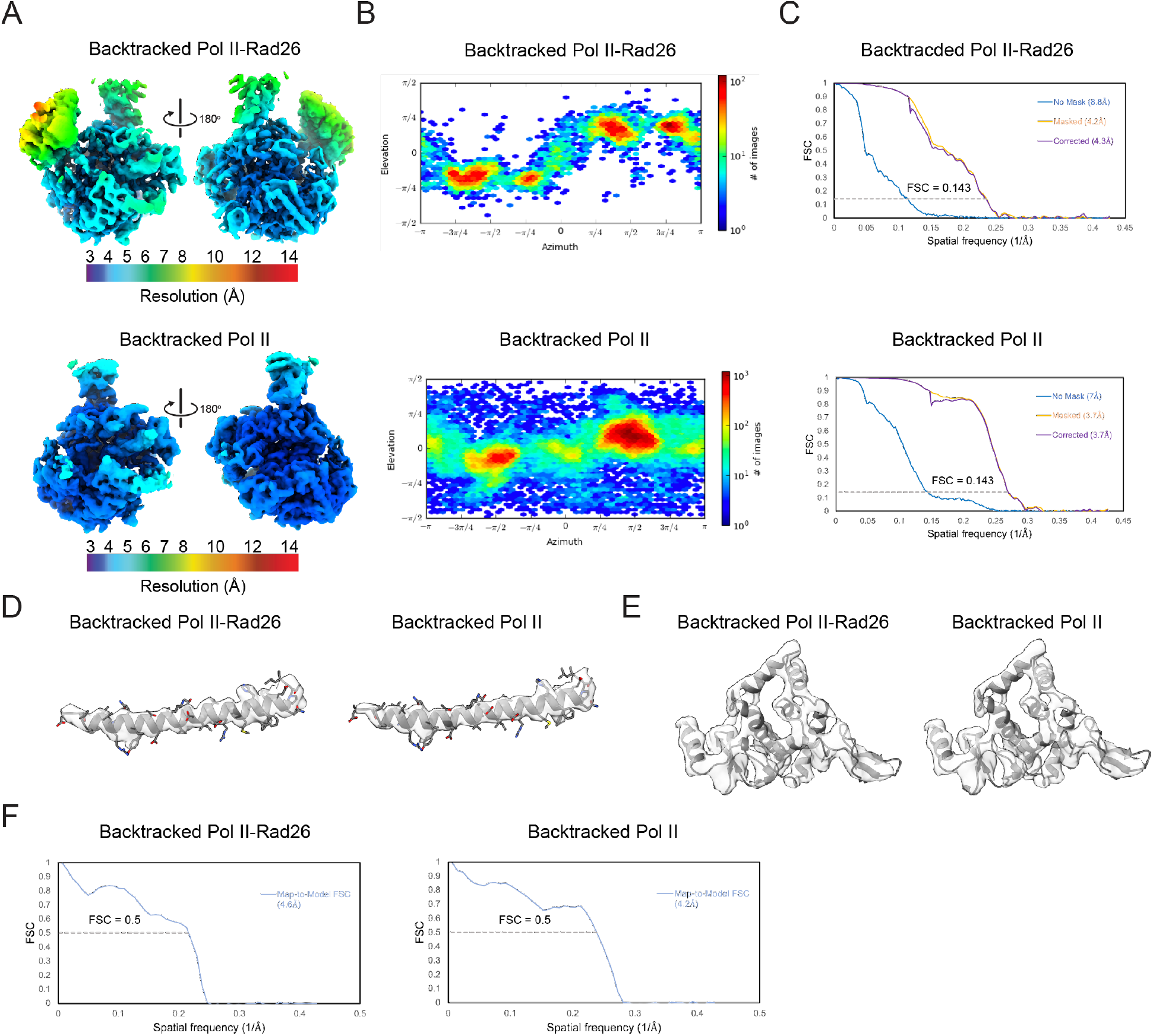
Analysis of the Backtracked Pol II-Rad26 and Backtracked Pol II cryo-EM maps. **(A)** Front and back views of locally filtered maps, colored by local resolution, of Backtracked Pol II-Rad26 and Backtracked Pol II. **(B, C)** Euler angle distribution of particle images **(B)** and FSC plots **(C)** for the maps shown in (A). **(D-E)** Close-ups of the cryo-EM densities corresponding to the Rpb1 Bridge helix **(D)**, and the Rpb2/Rpb9 ‘Jaw’ of Pol II **(E)** for the indicated structures with the models fitted in. **(F)** FSC curves for map-to-model fits for the maps shown in (A). The 0.5 FSC line is shown.

**Figure S5.**
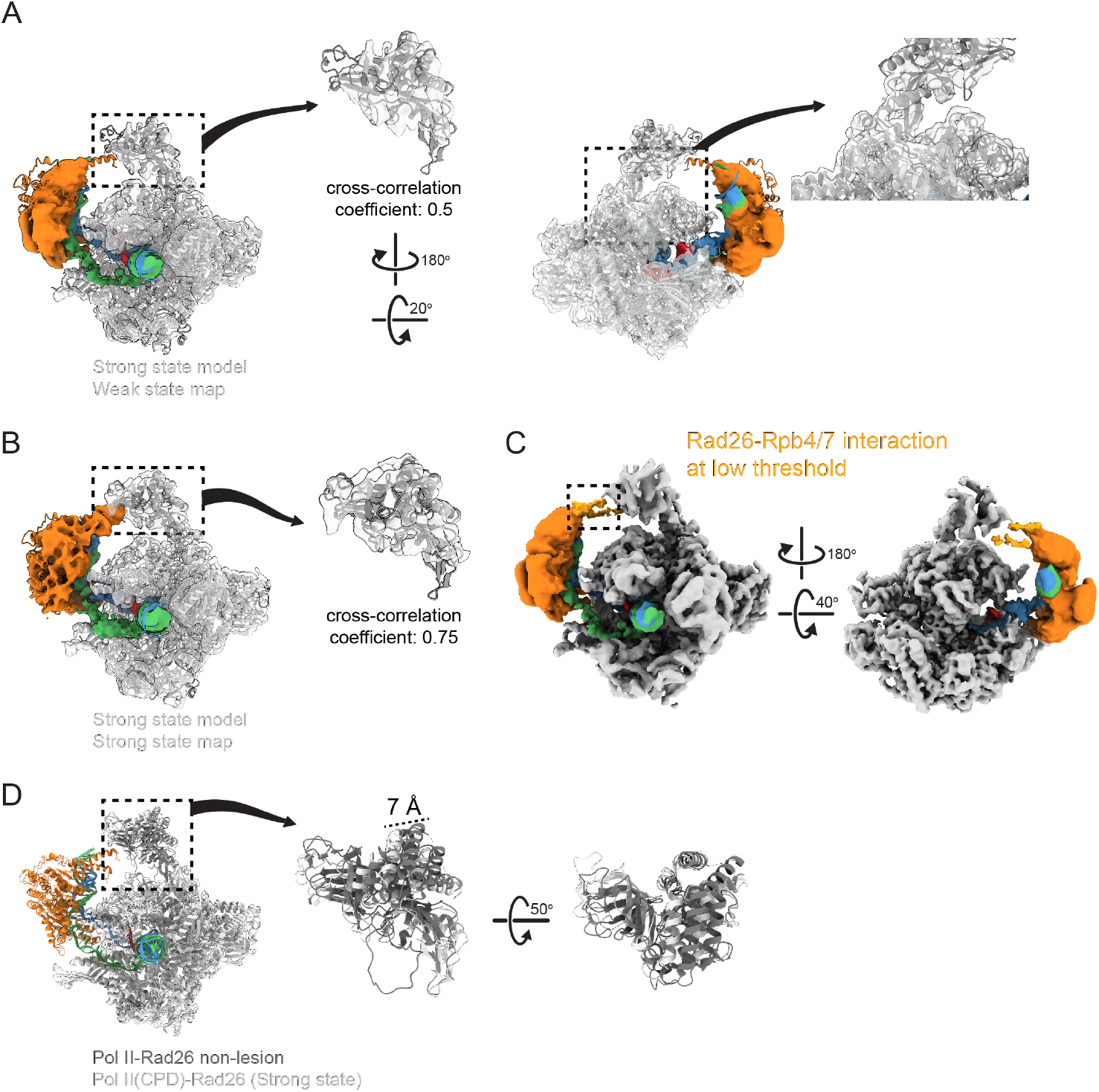
Structural analysis of Pol II-Rad26 and Pol II(CPD)-Rad26 complexes. **(A)** Two views are shown of the model for the Strong Rad26-Rpb4/7 state of Pol II(CPD)-Rad26 fitted into the cryo-EM map of the Weak Rad26-Rpb4/7 state of Pol II(CPD)-Rad26, with zoomed-in view of the cryo-EM density of Rpb4/7 with the model fitted in shown to their right. Fitting of the model into the map was driven by the core of Pol II. The cross-correlation coefficient for the fitting of the Rpb4/7 model for the Strong state into the map of the Weak state was 0.5 as reported by Fit-in-Map in ChimeraX. **(B)** Model for the Strong Rad26-Rpb4/7 state fitted into the cryo-EM map for the same state. The cross-correlation coefficient for the fitting of the Rpb4/7 model for the Strong state into the map of the Strong state was 0.75 as reported by Fit-in-Map in ChimeraX. **(C)** Cryo-EM map of the Weak Rad26-Rpb4/7 state of Pol II(CPD)-Rad26 shown at lower threshold, where the interaction between Rad26 and Rpb4/7 becomes apparent. **(D)** Superposition of models for Pol II(CPD)-Rad26 (Strong state) and Pol II-Rad26 (no lesion). The models were aligned using the core of Pol II. Two zoomed-in views of Rpb4/7 from the two models are shown to the right.

**Figure S6.**
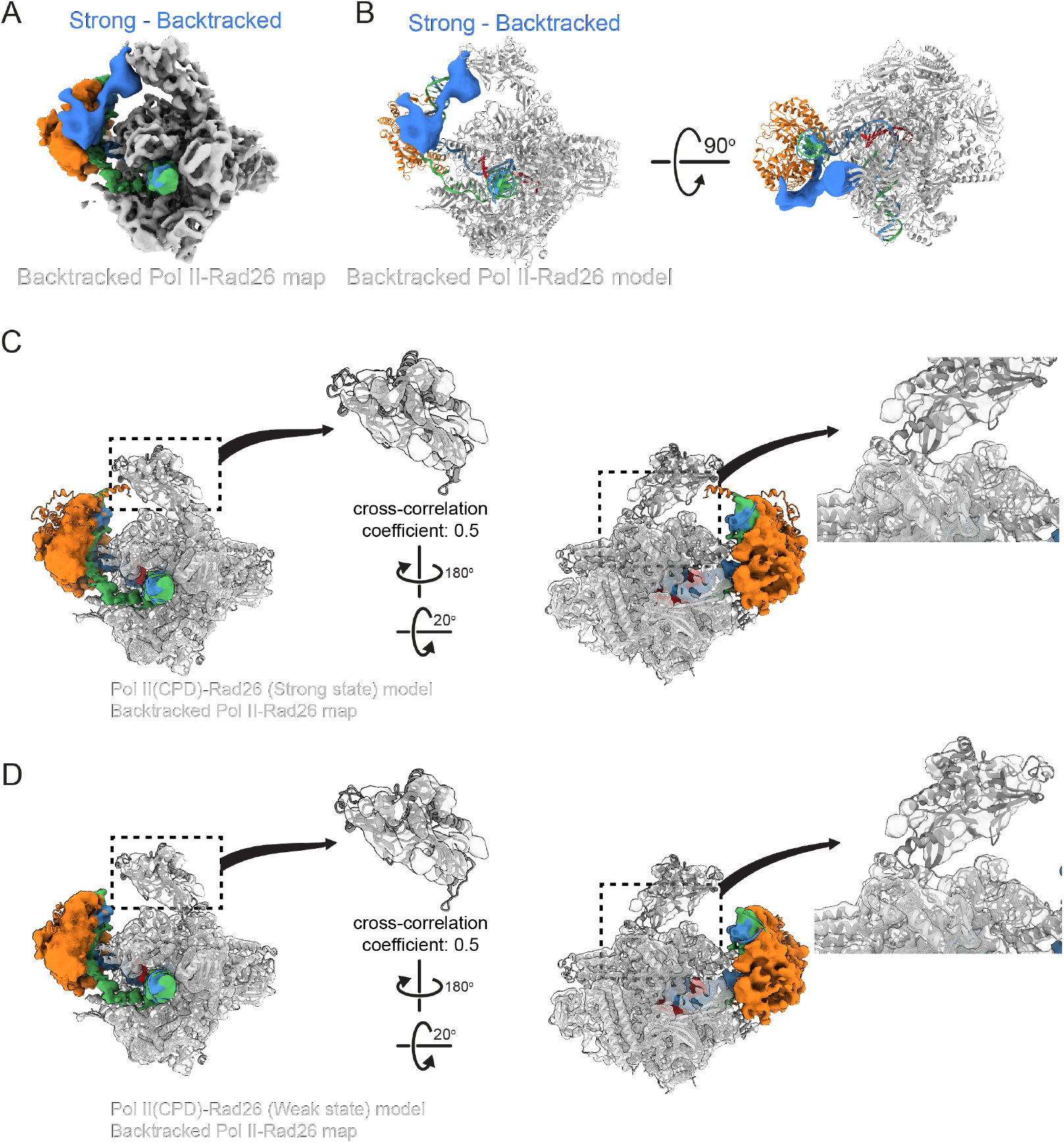
The Rad26-Rpb4/7 interaction is weakest in Backtracked Pol II-Rad26. **(A, B)** Difference map (in blue) calculated by subtracting Backtracked Pol II-Rad26 from Pol II(CPD)-Rad26 (Strong state), displayed on either **(A)** the cryo-EM density or **(B)** the atomic model for Backtracked Pol II-Rad26. **(C)** Two views are shown of the model for Pol II(CPD)-Rad26 (Strong state) fitted into the cryo-EM map of the Backtracked Pol II-Rad26, with zoomed-in views of the cryo-EM density of Rpb4/7 with the model fitted in shown to their right. The cross-correlation coefficient for the fitting of the Rpb4/7 model for the Strong state into the map of Backtracked Pol II-Rad26 was 0.5 as reported by Fit-in-Map in ChimeraX. **(D)** same as (C), but with the model for Pol II(CPD)-Rad26 (Weak state) fitted into the cryo-EM map of the Backtracked Pol II-Rad26. The cross-correlation coefficient for the fitting of the Rpb4/7 model for the Weak state into the map of Backtracked Pol II-Rad26 was 0.5 as reported by Fit-in-Map in ChimeraX.

**Figure S7.**
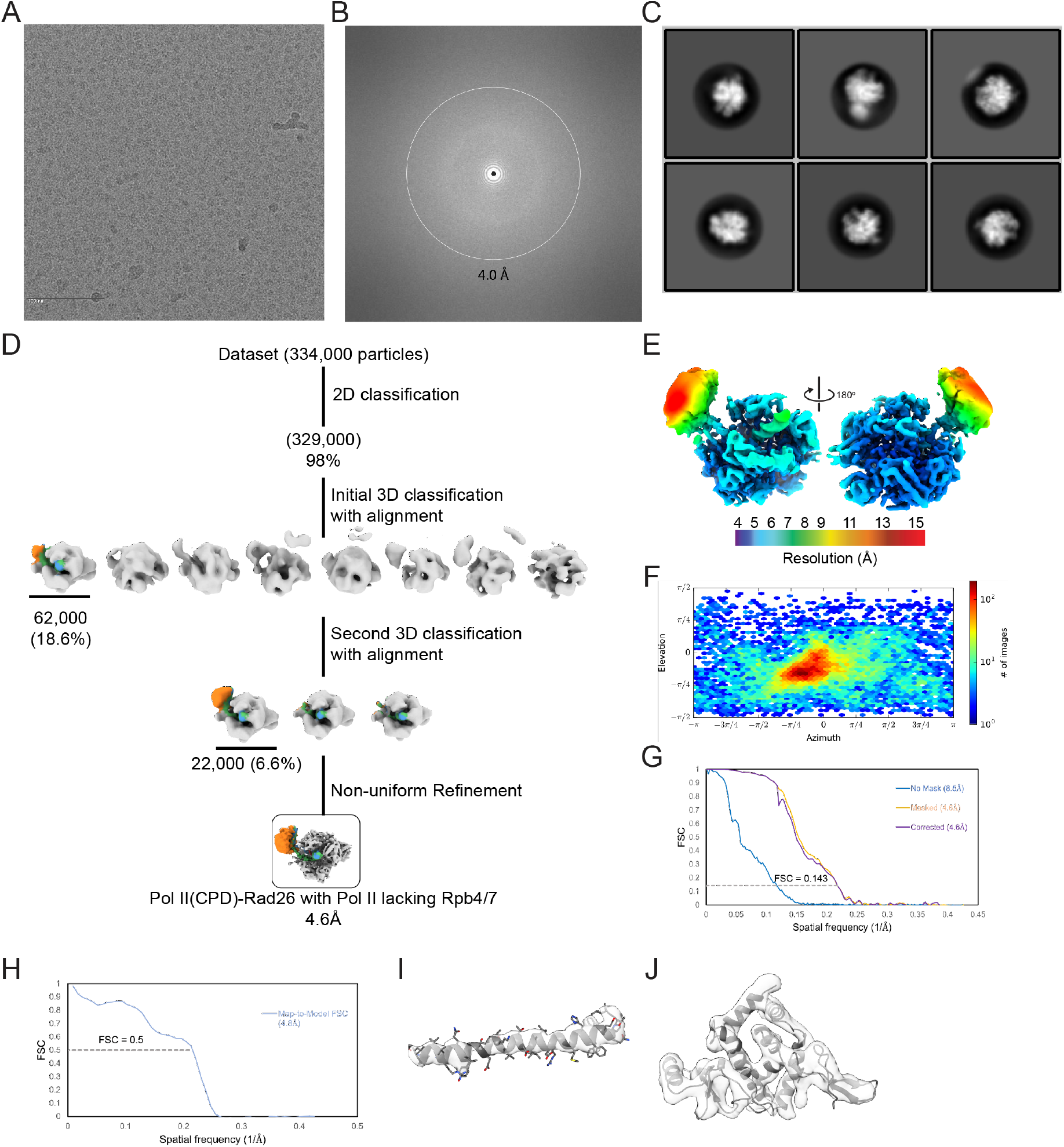
Cryo-EM structure determination and analysis of the Pol II(CPD)-Rad26 complex with Pol II lacking Rpb4/7. **(A-C)** Representative micrograph **(A)**, power spectrum **(B)**, and representative 2D class averages **(C)** of Pol II(CPD)-Rad26 with Pol II lacking Rpb4/7. **(D)** Schematic representation of the strategy used to sort out the complex particles. The number of particles contributing to each selected structure is indicated. The percentages shown are related to the total number of particles picked from micrographs. The indicated resolution corresponds to the 0.143 Fourier shell correlation (FSC) based on gold-standard FSC curves. **(E)** Front and back views of locally filtered maps, colored by local resolution, of Pol II(CPD)-Rad26 with Pol II lacking Rpb4/7. **(F-H)** Euler angle distribution of particle images **(F)**, FSC plot **(G)** and FSC curve for the map-to-model fit **(H)** for the map shown in (E). **(I**,**J)** Close-ups of the cryo-EM densities corresponding to the Rpb1 Bridge helix **(I)**, and the Rpb2/Rpb9 ‘Jaw’ of Pol II **(J)** for the indicated structure with the model fitted in.

**Figure S8.**
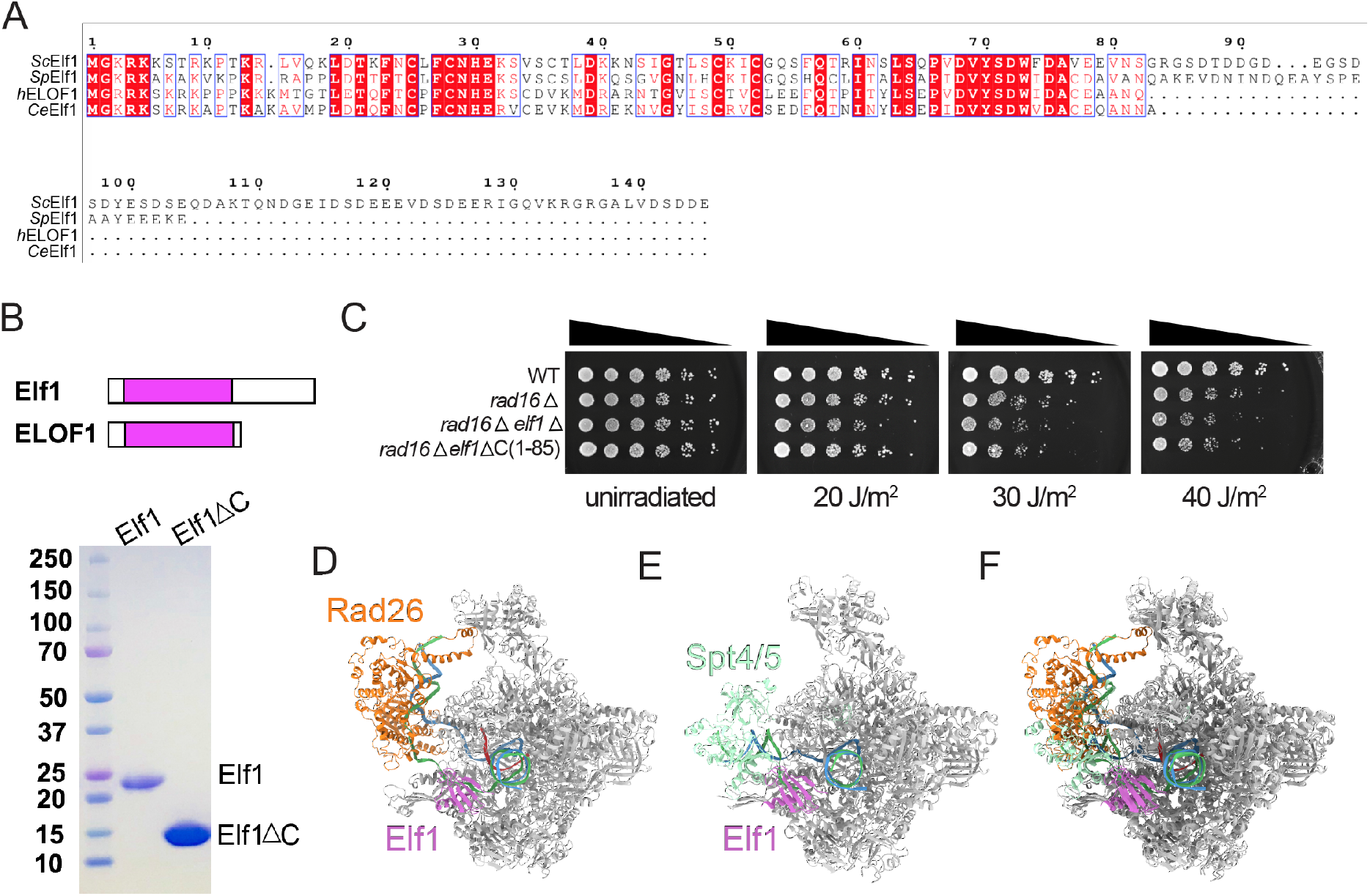
Purification of Elf1 and comparison of our structure of Pol II(CPD)-Rad26-Elf1 with the published structure of Pol II-Spt4/5-Elf1. **(A)** Sequence alignment of Elf1 orthologs from *S*.*cerevisiae* (*Sc*), *S*.*pombe* (*Sp*), humans (*h*) and *C. elegans* (*Ce*). **(B)** SDS-PAGE of purified Elf1 and Elf1core, shown schematically at the top. **(C-D)** Structures of **(C)** Pol II(CPD)-Rad26-Elf1 (this work) and **(D)** Pol II-Spt4/5-Elf1 (PDB: 6J4Y)(1). **(E)** Superimposition of the two models in (C) and (D).

**Figure S9.**
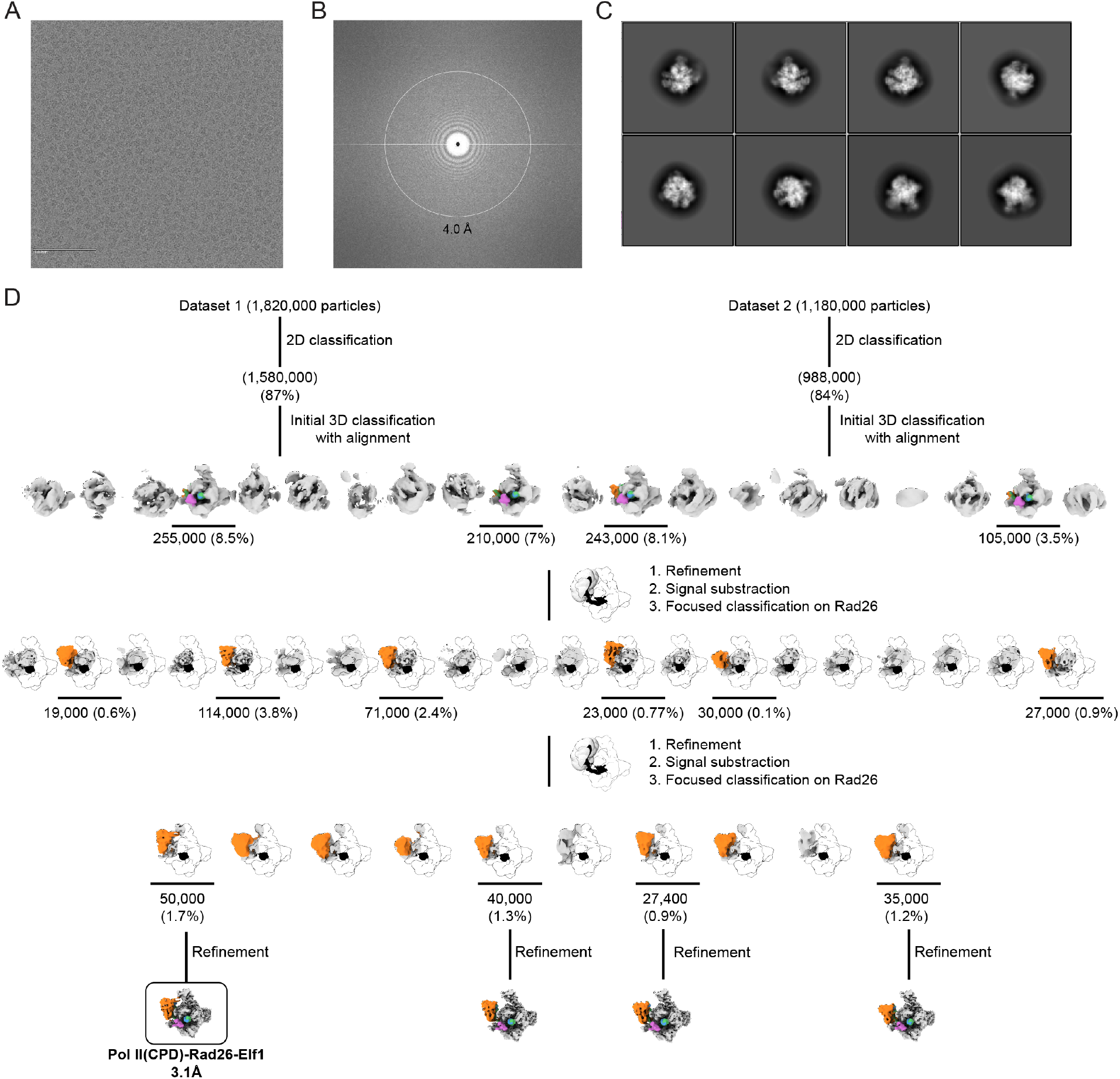
Cryo-EM structure determination of the Pol II(CPD)-Rad26-Elf1 complex. **(A-C)** Representative micrograph **(A)**, power spectrum **(B)**, and representative 2D class averages **(C)** of the Pol II(CPD)-Rad26-Elf1 complex. **(D)** Schematic of the strategy used to sort out the dataset. Focused 3D classification was performed without alignment unless otherwise noted. The number of particles contributing to each selected structure is indicated. The percentages shown are related to the total number of particles picked from the micrographs. The indicated resolution corresponds to the 0.143 Fourier shell correlation (FSC) based on gold-standard FSC curves (see Figure S10).

**Figure S10.**
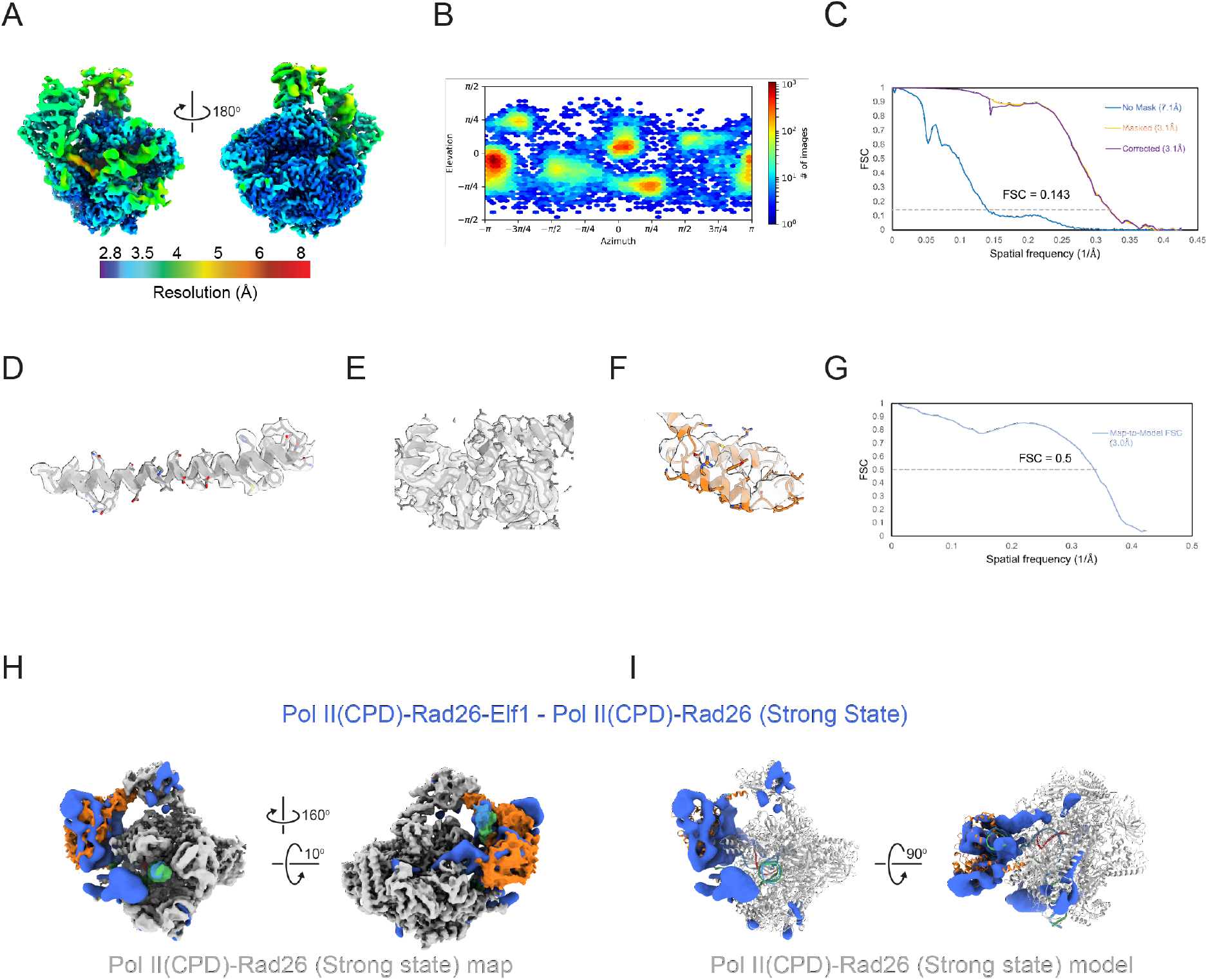
Analysis of the Pol II(CPD)Rad26-Elf1 cryo-EM map. **(A)** Front and back views of locally filtered maps, colored by local resolution. **(B, C)** Euler angle distribution of particle images **(B)** and FSC plots **(C)** for the map shown in (A). **(D-F)** Close-ups of the cryo-EM densities corresponding to the Rpb1 Bridge helix **(D)**, the Rpb2/Rpb9 ‘Jaw’ of Pol II **(E)** and the Rad26 HD2-1 ‘wedge’ **(F)** for the indicated structures with the models fitted in. **(G)** FSC curves for map-to-model fit for the map shown in (A). The 0.5 FSC line is shown. **(H, I)** Difference map (in blue) calculated by subtracting Pol II(CPD)Rad26 (Strong state) from Pol II(CPD)-Rad26-Elf1, displayed on either **(H)** the cryo-EM density or **(I)** the atomic model for Pol II-CDP-Rad26 (Strong state).

**Figure S11.**
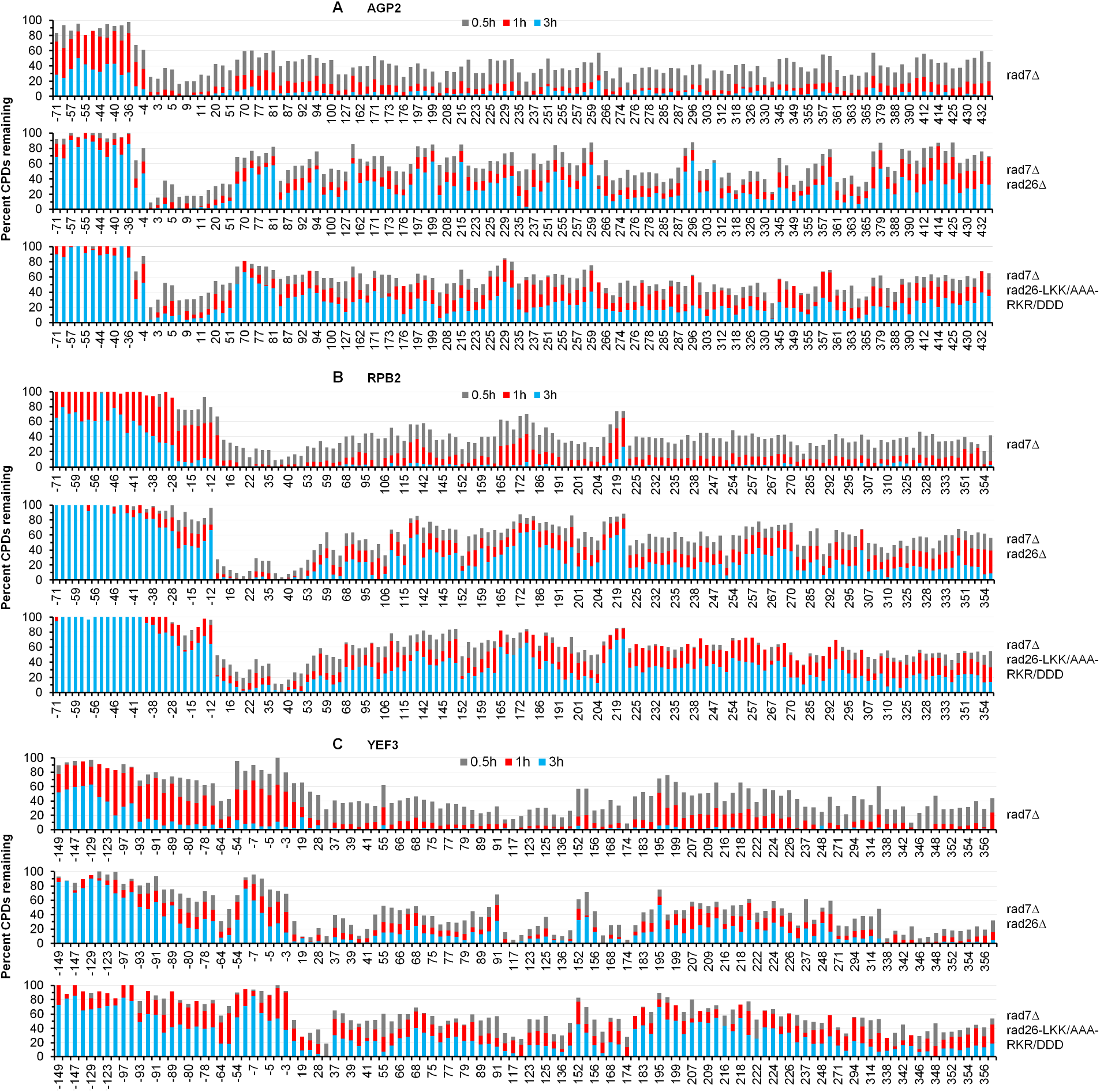
Base-resolution measurement of remaining CPD distributions at different loci after TC-NER. **(A-C)** Fraction (%) of CPDs at the indicated times of repair incubation along the AGP2 **(A)**, RPB2 **(B)** and YEF3 **(C)** loci of the indicated strains. Numbers at the bottom of each plot indicating the nucleotide positions of the loci are relative to the major TSS (+1).

**Table S1.** Cryo-EM data collection, refinement and validation statistics.

|  | Pol II(CPD)-Rad26 Dataset |  |  |  | 10-subunit |
| --- | --- | --- | --- | --- | --- |
|  | Strong state | Weak state | Pol II(CD) Conf1 | Pol II(CPD) Conf2 |  |
| **PDB** | 8TUG | 8TVP | 8TVW | 8TVX | 8TVQ |
| **EMDB** | 41623 | 41647 | 41653 | 41654 | 41648 |
| **Data Collection** |  |  |  |  |  |
| Microscope |  |  | Talos Arctica |  | Talos Arctica |
| Camera |  |  | K2 Summit |  | K2 Summit |
| Camera Mode |  |  | Counting |  | Super-Res |
| Voltage (kV) |  |  | 200 |  | 200 |
| Magnification |  |  | 36,000 |  | 36,000 |
| Pixel Size (Å/pixel) |  |  | 1.16 |  | 1.16 |
| Dose rate (e-/Å<sup>2</sup> second) |  |  | 8.4 |  | 4 |
| Total dose (e-/Å<sup>2</sup>) |  |  | 59 |  | 52 |
| Number of frames |  |  | 47 |  | 52 |
| Defocus range (µm) |  |  | 0.6-2.5 |  | 0.6-2.5 |
| Micrographs collected (no.) |  |  | 3,358 |  | 955 |
| Initial particle (no.) |  |  | 1,620,000 |  | 334,000 |
| Final particle (no.) | 20,000 | 25,000 | 74,000 | 73,000 | 22,000 |
| **Refinement** |  |  |  |  |  |
| Initial model used |  |  | 1Y77 |  | 1Y77 |
| Final resolution (Å) | 3.5 | 3.7 | 3.6 | 3.7 | 4.6 |
| (0.143 FSC threshold) |  |  |  |  |  |
| Map sharpening B factor (Å<sup>2</sup>) | -73 | -68 | -87 | -101 | -153 |
| **Model Refinement** |  |  |  |  |  |
| Map-to-model resolution (Å) | 3.6 | 3.8 | 3.8 | 3.9 | 4.8 |
| (0.5 FSC threshold) |  |  |  |  |  |
| *Model Composition* |  |  |  |  |  |
| Nonhydrogen atoms | 71,184 | 69,002 | 61,459 | 60,948 | 64,987 |
| Protein residues | 4,251 | 4,182 | 3,748 | 3,748 | 3,869 |
| Nucleotides | 103 | 103 | 56 | 56 | 100 |
| Ligands | 9 | 9 | 9 | 9 | 9 |
| B factor (Å<sup>2</sup>) | 244 | 205 | 123 | 170 | 283 |
| *R.m.s. deviations* |  |  |  |  |  |
| Bond length (Å) | 0.003 | 0.003 | 0.003 | 0.003 | 0.003 |
| Bond angle (°) | 0.56 | 0.54 | 0.59 | 0.60 | 0.644 |
| *Validation* |  |  |  |  |  |
| MolProbity score | 1.7 | 1.8 | 1.7 | 1.7 | 2.1 |
| Clash score | 6.7 | 8.6 | 7.0 | 9.2 | 14.9 |
| Poor rotamers (%) | 0 | 0 | 0 | 0 | 0 |
| *Ramachandran* |  |  |  |  |  |
| Favored (%) | 95.53 | 95.60 | 96.0 | 95.82 | 93.79 |
| Allowed (%) | 4.4 | 4.3 | 4.0 | 4.1 | 6.1 |
| Disfavored (%) | 0.07 | 0.1 | 0.05 | 0.08 | 0.11 |

Table S1. Cryo-EM data collection, refinement and validation statistics (continued)
|  | **Backtracked Pol II-Rad26 Dataset** |  | **Pol II(CPD) Rad26-Elf1** |
| --- | --- | --- | --- |
|  | **Backtracked Pol II-Rad26** | **Backtracked Pol II** |  |
| **PDB** | 8TVS | 8TVV | 8TVY |
| **EMDB** | 41650 | 41652 | 41655 |
| **Data Collection** |  |  |  |
| Microscope | Talos Arctica |  | Talos Arctica |
| Camera | K2 Summit |  | K2 Summit |
| Camera Mode | Counting |  | Counting |
| Voltage (kV) | 200 |  | 200 |
| Magnification | 36,000 |  | 36,000 |
| Pixel Size (Å/pixel) | 1.16 |  | 1.16 |
| Dose rate (e-/Å<sup>2</sup> second) | 5 |  | 5/5.5 |
| Total dose (e-/Å<sup>2</sup>) | 55 |  | 50/55 |
| Number of frames | 55 |  | 50 |
| Defocus range (µm) | 0.6-2.5 |  | 0.6-2.5 |
| Micrographs collected (no.) | 9,167 |  | 8,000 |
| Initial particle (no.) | 3,310,000 |  | 3,000,000 |
| Final particle (no.) | 11,000 | 100,000 | 50,000 |
| **Refinement** |  |  |  |
| Initial model used | 1Y77 | 1Y77 | 1Y77 |
| Final resolution (Å)<br>(0.143 FSC threshold) | 4.4 | 3.7 | 3.1 |
| Map sharpening *B* factor (Å<sup>2</sup>) | -92 | -117 | -85 |
| **Model Refinement** |  |  |  |
| Map-to-model resolution (Å)<br>(0.5 FSC threshold) | 4.6 | 4.2 | 3.0 |
| *Model Composition* |  |  |  |
| Nonhydrogen atoms | 70,048 | 61,419 | 77,243 |
| Protein residues | 4,182 | 3,747 | 4,701 |
| Nucleotides | 104 | 56 | 103 |
| Ligands | 9 | 9 | 9 |
| *B* factor (Å<sup>2</sup>) | 266 | 149 | 89 |
| *R.m.s. deviations* |  |  |  |
| Bond length (Å) | 0.003 | 0.003 | 0.003 |
| Bond angle (°) | 0.553 | 0.555 | 0.594 |
| *Validation* |  |  |  |
| MolProbity score | 1.8 | 1.7 | 1.9 |
| Clash score | 7.9 | 8.1 | 7.9 |
| Poor rotamers (%) | 0 | 0 | 0 |
| *Ramachandran* |  |  |  |
| Favored (%) | 95.36 | 95.88 | 92.49 |
| Allowed (%) | 4.52 | 4.07 | 7.45 |

**Table S2.**
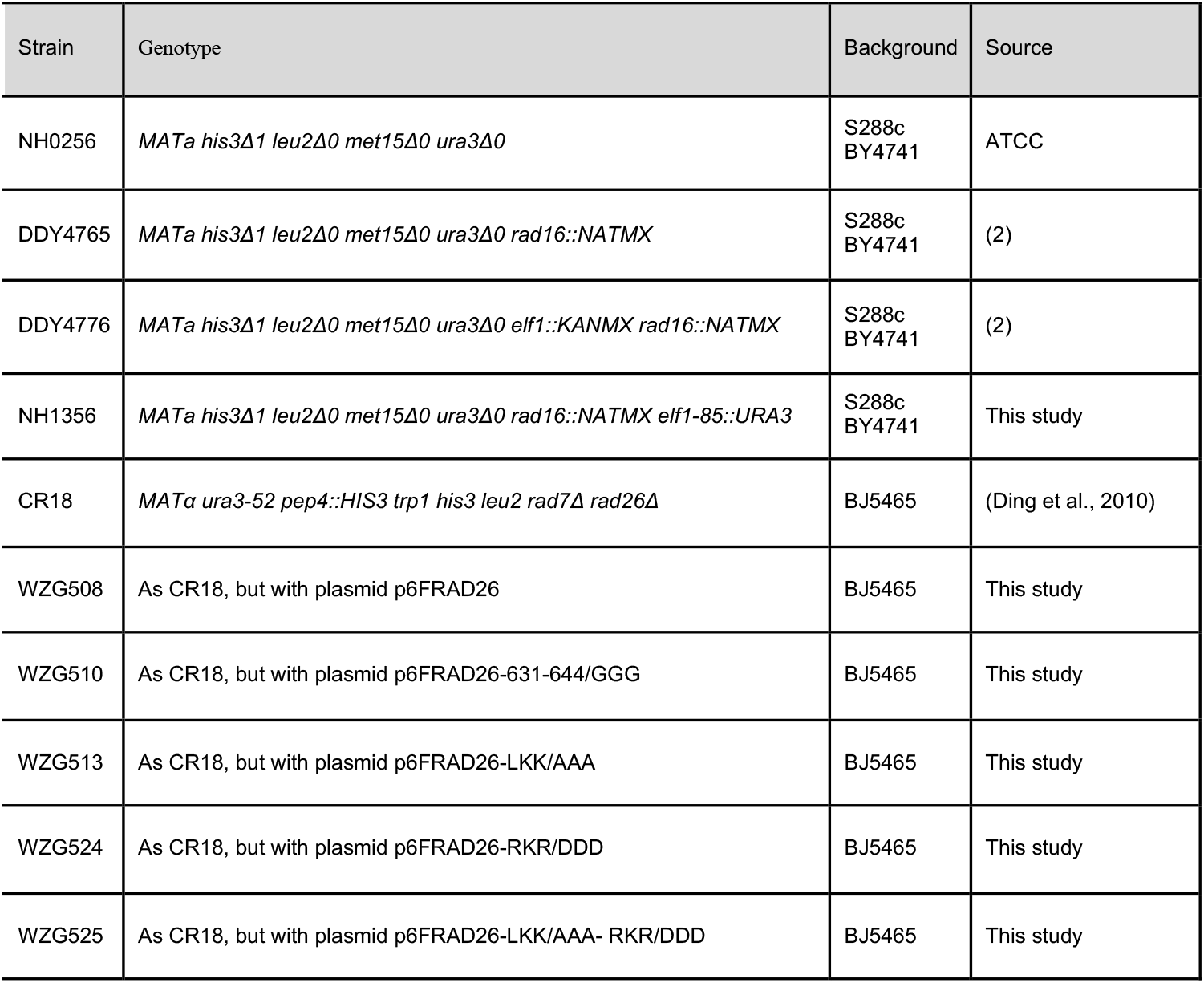
Saccharomyces cerevisiae strains.

